# A fungal plant pathogen overcomes *mlo*-mediated broad-spectrum disease resistance by rapid gene loss

**DOI:** 10.1101/2021.12.09.471931

**Authors:** Stefan Kusch, Lamprinos Frantzeskakis, Birthe D. Lassen, Florian Kümmel, Lina Pesch, Mirna Barsoum, Kim D. Walden, Ralph Panstruga

## Abstract

Hosts and pathogens typically engage in a co-evolutionary arms race. This also applies to phytopathogenic powdery mildew fungi, which can rapidly overcome plant resistance and perform host jumps. Using experimental evolution, we show that the powdery mildew pathogen *Blumeria hordei* is capable of breaking the agriculturally important broad-spectrum resistance conditioned by barley loss-of-function *mlo* mutants. Partial *mlo* virulence of evolved *B. hordei* isolates is correlated with a distinctive pattern of adaptive mutations, including small-sized (8-40 kb) deletions, of which one is linked to the *de novo* insertion of a transposable element. Occurrence of the mutations is associated with a transcriptional induction of effector protein-encoding genes that is absent in *mlo*-avirulent isolates on *mlo* mutant plants. The detected mutational spectrum comprises the same loci in at least two independently isolated *mlo*-virulent isolates, indicating convergent multigenic evolution. The mutational events emerged in part early (within the first five asexual generations) during experimental evolution, likely generating a founder population in which incipient *mlo* virulence was later stabilized by additional events. This work highlights the rapid dynamic genome evolution of an obligate biotrophic plant pathogen with a transposon-enriched genome.

## Introduction

Pathogens and their hosts are locked in a coevolutionary competition where the host attempts to prevent pathogen infection, while the pathogen adapts to evade host recognition and retain its ability to infect the host. Generalist pathogens infect a broad range of hosts, and their genomes evolve in response to selection by many hosts under diffuse coevolution [21]. Specialist pathogens, on the other hand, infect one or few hosts and are engaged in an intimate coevolutionary arms race. The gene-for-gene hypothesis is a paradigm for the arms race between plants and their pathogens [24]. Plants have intracellular nucleotide-binding oligomerization domain (NOD)-like receptors (NLRs) that can recognize pathogen effectors and mount an effective defense response, while pathogens evolve to subvert or circumvent perception [40]. Obligate biotrophic pathogens depend on living host cells and, thus, have to evade recognition by NLRs for their very survival and reproduction. Such specialist pathogens often rapidly overcome resistance conferred by effector recognition via cognate host NLRs. For example, the obligate biotrophic barley powdery mildew fungus *Blumeria hordei* (*B. hordei*) frequently escapes resistance conditioned by alleles of *Mildew locus A* (*Mla*)-encoded NLRs in barley (*Hordeum vulgare*) [11]. Mla immune receptors directly bind secreted *B. hordei* effectors, and loss of recognition happens by loss or modification of the cognate effector due to spontaneous mutations in the fungal genome [58, 80]. In addition, copy number variation of effector genes contributes to genetic variation and can lead to outbreaks of new strains of filamentous plant pathogens [72, 36].

Short generation times, large effective population sizes, and plastic genome architectures are drivers of rapid adaptation in microbial pathogens [26]. Within the lifetime of a plant, pathogens can go through tens or even hundreds of generation cycles, and can produce thousands to millions of individual mito- and meiospores as offspring [88]. Hence, the standing genetic variation of the pathogen population is much larger than that of its host by default. This is likely a prerequisite for pathogen survival, since recognition of a single effector suffices to prevent proliferation and reproduction.

The genomes of filamentous plant pathogens frequently exhibit special architectures and features including dispensable accessory chromosomes and hypervariable minichromosomes [50]. These are often enriched with transposable elements, carry effector genes, and are otherwise gene-poor. Binary genome compartmentalization, known as two-speed architecture, is characterized by distinct transposable element-rich regions that mainly contain effector genes that evolve at a faster pace than the rest of the genome [20, 74]. Intriguingly, some fungal plant pathogens including the cereal powdery mildew pathogen *Blumeria* harbor genomes that are massively inflated by transposable elements [27, 63, 64]. Transposable elements are equally distributed throughout these genomes, which are further characterized by extensive copy number variation of effector genes and the loss of some conserved ascomycete genes [27, 85]. In particular, long interspersed nuclear element (LINE) and long terminal repeat (LTR) retrotransposons are highly abundant, and these elements exhibit mostly very low sequence divergence, suggesting that recent transposon bursts shaped the genome of *B. hordei* [27].

At least 900 species of powdery mildew fungi (Ascomycota, *Erysiphaceae*) infect >10,000 plant species worldwide [10, 49], including crops, trees, and herbs [31]. The pathogen causes significant yield losses if not held in check by fungicides, affecting grain yield and quality [18]. Even though powdery mildews apparently occur as homogenous strains due to their dominating asexual mode of propagation, they can have a complex population structure with many different haplotypes present within a supposedly clonal isolate [6].

The loss-of-function mutation of *Mildew resistance Locus O* (*MLO*) gene(s) confers highly effective and durable broad-spectrum resistance against powdery mildew in many plant species [47]. Pathogenesis is terminated prior to fungal host cell entry on *mlo* mutants, which have been used widely in European barley agriculture since the late 1970s [41]. *MLO* genes encode seven-transmembrane domain proteins [19, 12] with a cytosolic calmodulin-binding domain in the carboxyterminus [44] and function as cation channels [29]. Genetic suppressors of *mlo*-based resistance cause partial susceptibility to powdery mildew. These include the *Required for mlo-specified resistance* (*Ror*) genes in barley [28, 16, 1], as well as *PENETRATION* (*PEN*) genes in *Arabidopsis thaliana*, which are major components of pre-penetration resistance to powdery mildew [16, 55, 87, 38]. In addition, abiotic stress conditions can result in a temporary break-down of *mlo*-based resistance [81, 65]. The Japanese *B. hordei* isolate RACE1 is the only known natural case with partial virulence on barley *mlo* mutant plants [59, 42]. In addition, a previous experimental evolution approach revealed several partially *mlo*-virulent *B. hordei* isolates [82]. Molecular analysis of these *B. hordei* isolates indicated that the *mlo* virulence phenotype depends on a small number of unidentified genes [33, 3], but the underlying mechanism remains elusive.

To close this gap in knowledge, we here deployed realtime evolution experiments to select a set of *mlo*-virulent *B. hordei* isolates for detailed molecular analysis. We identified a distinctive pattern of few convergent mutational events in the three identified isolates, resulting in an altered transcriptional program during fungal pathogenesis on barley *mlo* mutant plants. Our data suggest that the majority of mutational events occurred early in our experimental setup and rapidly outcompeted the respective parental alleles. Enhanced *B. hordei* virulence on *mlo* mutants seems to be correlated with a fitness cost in the form of lowered virulence on barley wild-type plants, which may explain why the *mlo*-virulent phenotype does not prevail under agricultural conditions. Collectively, our findings provide an example of how few mutations in the genome of a phytopathogen can cause a drastic change in its virulence spectrum, enabling its rapid adaptation to new host environments.

## Results

### Selection of three partially *mlo*-virulent *B. hordei* isolates by experimental evolution

To study the rapid evolution of *B. hordei* experimentally, we selected for *mlo*-virulent *B. hordei* isolates derived from the *mlo*-avirulent parental strain *B. hordei* K1_AC_. For 15 asexual generations, occasionally occurring *B. hordei* K1_AC_ colonies on otherwise highly resistant barley *mlo* mutant plants were recovered, fungal biomass proliferated on the susceptible (*Mlo* genotype) cultivar (cv.) Ingrid, and then conidia re-inoculated on *mlo* plants, as described before [82]. Subsequently, from generation 15 onwards, the resulting *B. hordei* isolates were cultivated on barley *mlo* mutant plants only, yielding the three independent strains Supervirulent K1_AC_ (SK1), SK2, and SK3. These isolates showed stable yet partial *mlo* virulence with visible sporulation and a 15-25% host cell entry rate on otherwise highly resistant barley *mlo*-3 backcrossed to cv. Ingrid (BCI) mutant plants (Figure 1A and 1B). *B. hordei* SK1 exhibited a similar level of virulence on a range of near-isogenic BCI *mlo* lines with various mutational defects in *Mlo* [75], indicating that in contrast to *B. hordei* RACE1 [95] the enhanced virulence of this strain is independent of the host *mlo* allele (Figure 1C and Supplementary Figure 1). We found comparable host cell entry levels on two *mlo* mutants in different barley lines, Pallas *mlo*-5 (approx. 15%) and Haisa *mlo*-1 (approx. 20%; Figure 1D), demonstrating that the *B. hordei* SK1 virulence phenotype is also independent of the host genetic background. We nevertheless noticed a slight but statistically significant reduction of *B. hordei* SK1 host cell entry rates compared to *B. hordei* K1_AC_ on all wild-type (*Mlo*) genotypes tested (cv. Ingrid, Pallas and Haisa; Figure 1B and 1C), suggesting a reduction in pathogenic fitness in the *mlo*-virulent fungal isolate (see also below).

**Fig. 1.**
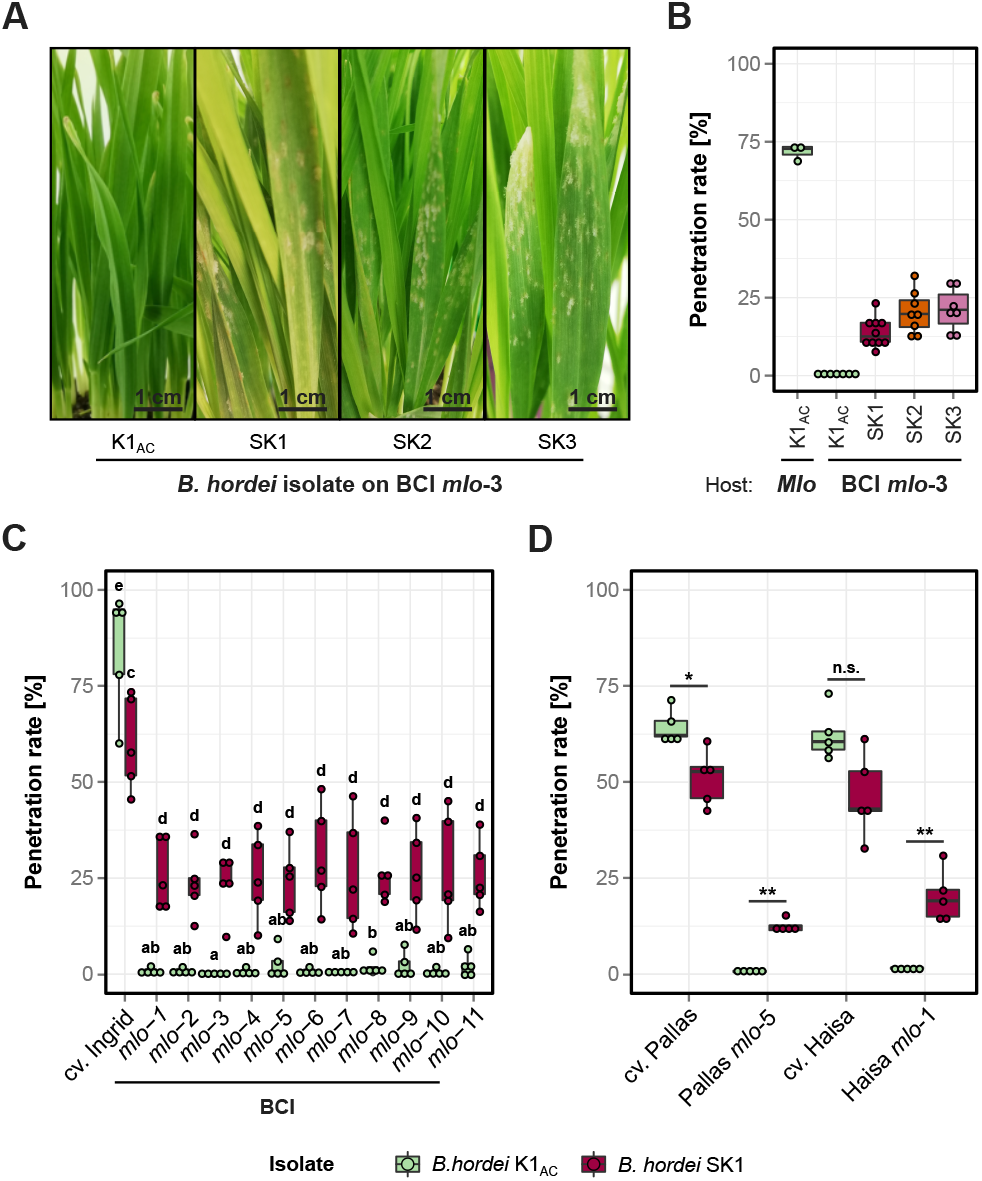
Isolation of three partially *mlo*-virulent *B. hordei* isolates. (**A**) Powdery mildew symptoms caused by *B. hordei* K1_AC_ and *B. hordei* SK1, SK2, and SK3 at seven days post-inoculation (dpi) on barley backcross Ingrid (BCI) *mlo*-3 plants. (**B**) Penetration success in percent [%] of conidia from *B. hordei* K1_AC_ (green) and the isolates *B. hordei* SK1 (maroon), SK2 (orange), and SK3 (purple) on the sus-ceptible barley cv. Ingrid (*Mlo*) or the BCI *mlo*-3 mutant lines. All data points are displayed. (**C**) Penetration success [%] of conidia from *B. hordei* K1_AC_ (green) and SK1 (maroon) on the barley cultivar Ingrid and 11 *mlo* mutant alleles backcrossed to Ingrid (BCI). (**D**) Penetration success [%] of conidia from *B. hordei* K1_AC_ (green) and SK1 (maroon) on barley *mlo*-5 and *mlo*-1 and the respective parental cultivars, cv. Pallas and Haisa. Data are based on *n* = 5 independent replicates (**C** and **D)**; at least 100 interactions per leaf and three leaves were scored for each replicate. Letters denote statistical groups at *P* < 0.05 (**C**) according to Kruskal-Willis and Mann-Whitney-Wilcoxon testing. (**D**) *, *P* < 0.05; **, *P* < 0.01; n.s., not significant.

### Enhanced virulence of *B. hordei* SK1 is restricted to *mlo* mutants in barley and *Arabidopsis thaliana*

To assess whether *B. hordei* SK1 has a generally altered virulence spectrum, we inoculated various host (barley) and nonhost (*A. thaliana*) genotypes with conidia of this isolate and assessed the infection success. On barley BCI *mlo ror1* and *mlo ror2* double mutants, which have partially compromised *mlo* resistance due to second-site mutations in the genes *Ror1* and *Ror2* [28], the *mlo*-virulent isolate *B. hordei* SK1 showed elevated entry rates compared to *mlo* single mutant plants, indicative of an additive effect of the host *ror* mutations and the presumed genomic alterations in the fungal pathogen (Supplementary Figure 2). On barley lines carrying *Mildew resistance locus A* (*Mla*) alleles conferring isolate-specific immunity with different levels of effectiveness against *B. hordei* K1 [42, 83], *B. hordei* SK1 showed reduced entry success, but also on the near-isogenic control cultivars lacking the respective *Mla* genes (Supplementary Figure 3), suggesting that the isolate is incapable of overcoming race-specific resistance. *B. hordei* SK1 was further unable to colonize eight wheat cultivars and exhibited entry success levels comparable to *B. hordei* K1_AC_, except on three cultivars where we observed a slight increase (Supplementary Figure 4A and 4B). Notably, *B. hordei* SK1 had significantly reduced penetration success on the *A. thaliana* mutants *pen2 pad4 sag101* and *pen1*, which are partially defective in resistance to the non-adapted *B. hordei* pathogen [16, 55], but increased entry success on the *A. thaliana mlo2* single mutant and the *mlo2 mlo6 mlo12 pen1 pen2* quintuple mutant. *B. hordei* SK1 even succeeded with occasional entry on the otherwise extremely resistant *mlo2 mlo6 mlo12* triple mutant, providing an additional link to *mlo* virulence also in the *B. hordei* nonhost species *A. thaliana* (Supplementary Figure 4C).

### *B. hordei* SK1 has an altered transcriptional profile during haustorium formation

To identify genes contributing to *mlo* virulence in *B. hordei* SK1 compared to *B. hordei* K1_AC_ we performed whole transcriptome shotgun sequencing (RNA-seq) at 6 hours post inoculation (hpi; around appressorium formation) and 18 hpi (around haustorium establishment in compatible interactions; Supplementary Table 1) using RNA extracted from inoculated barley (BCI *mlo*-3 genotype) leaf epidermal strips. The expression profiles of the host, representing the majority of the RNA-seq reads (>90%), did not vary significantly between the barley epidermal samples inoculated with the two *B. hordei* isolates according to principal component analysis (PCA) and hierarchical distance calculations (hierarchical clustering and non-metric dimensional scaling or NMDS; Figure 2A and Supplementary Figure 5). However, PCA revealed a clear separation of the host responses at 6 hpi and 18 hpi, i.e., by time, but not by fungal isolate (K1_AC_ versus SK1, Figure 2A). There were no differentially expressed (DE) host genes (inoculated by SK1 versus K1_AC_) at 6 hpi and nine DE host genes at 18 hpi (logFC > |1|, *P*_adj_ < 0.05; Supplementary Figure 6, Supplementary Table 2, Supplementary Table 3, Supplementary Table 4).

**Fig. 2.**
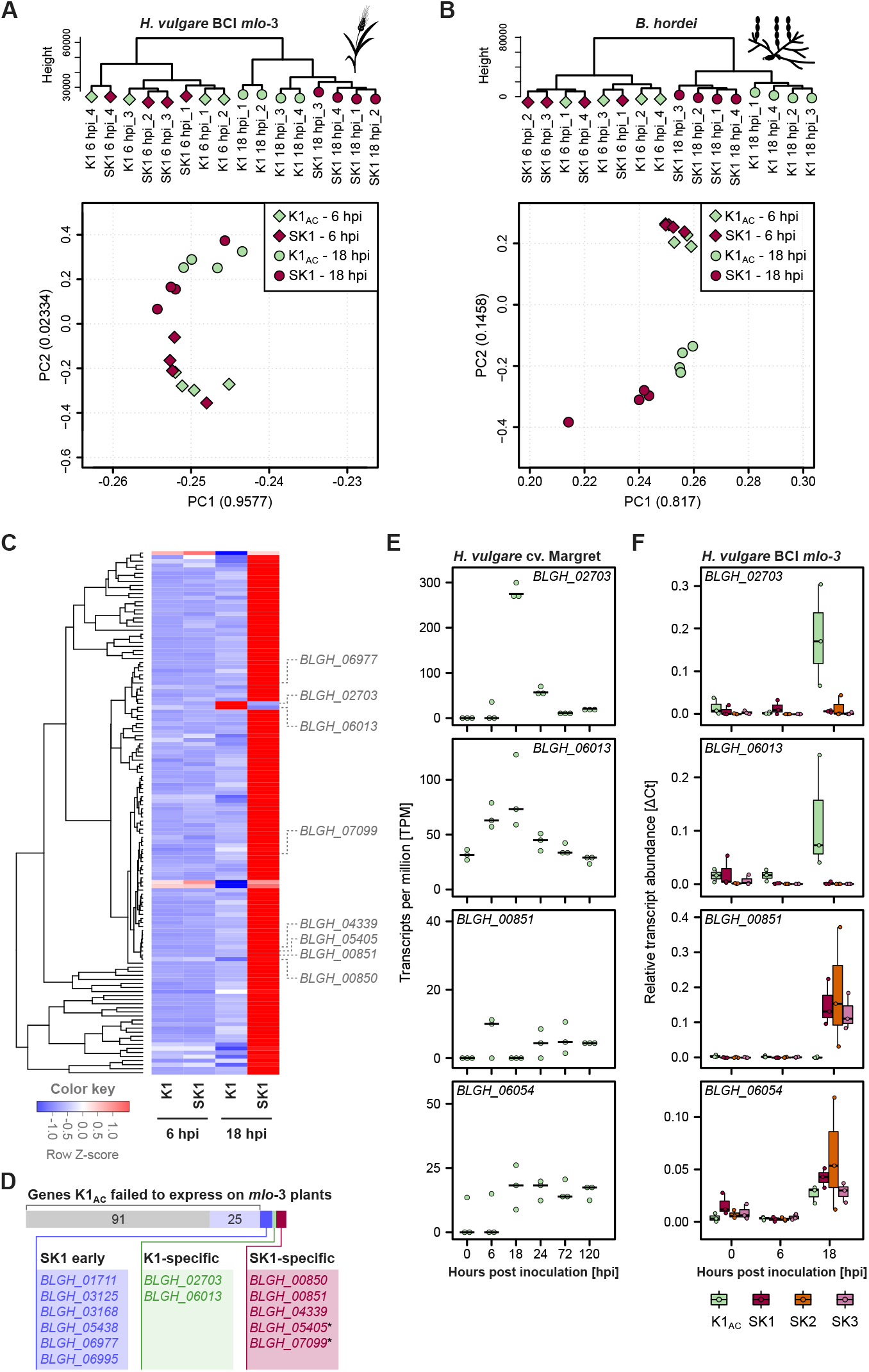
A distinct transcriptomic pattern is associated with *mlo* virulence of *B. hordei* isolate SK1. We conducted RNA-seq of epidermal samples collected from barley BCI *mlo*-3 inoculated with *B. hordei* K1_AC_ and SK1, respectively, at 6 hpi and 18 hpi, each with *n* = 4 independent replicates. (**A**) Hierarchical clustering dendrogram (upper panel) and principal component (PC) analysis (lower panel) of gene expression in barley BCI *mlo*-3 in the 18 samples accounting for the four conditions. Green, K1; maroon, SK1; squares, 6 hpi; circles, 18 hpi. Numbers in brackets indicate the ratio of data explained by the principal component. (**B**) Hierarchical clustering dendrogram and PCA of gene expression in *B. hordei* K1_AC_ and *B. hordei* SK1 at 6 hpi and 18 hpi on barley BCI *mlo*-3. (**C**) Differential expression analysis revealed 127 up-regulated and 2 down-regulated genes in *B. hordei* SK1 at 18 hpi on barley BCI *mlo*-3 (Supplementary Figure 6 and 7, Supplementary Table 6). The heat map shows the normalized relative expression (expressed as Row Z score) of these 129 genes in the two isolates at 6 and 18 hpi according to the color-coded scale at the bottom. Genes of particular interest within the heat map are indicated. (**D**) The illustration summarizes how the 129 DE SK1 genes compare with time course gene expression data obtained for K1_AC_ on the susceptible barley cv. Margret [70]. Grey indicates genes that are induced in SK1 at 18 hpi on cv. Margret but not expressed in K1_AC_ on barley *mlo*-3; light blue indicates genes down-regulated in K1_AC_ at 18 hpi compared to 6 hpi; dark blue labels genes highly up-regulated in SK1 at 18 hpi and up-regulated at later time points (24 hpi or after) in K1_AC_ on cv. Margret; green indicates genes down-regulated in SK1, which are up-regulated in K1_AC_ on cv. Margret at 18 hpi; red highlights genes specifically expressed in SK1. The asterisk indicates genes that were induced in K1_AC_ on cv. Margret, but only at marginal levels <50 TPM (Supplementary Figure S7). (**E**) Dot plots show the expression of *BLGH_02703, BLGH_06013, BLGH_00851*, and *BLGH_06054* in transcripts per million (TPM; y-axis) in *B. hordei* K1_AC_ on the susceptible cv. Margret in a time-course at 0, 6, 18, 24, 72, and 120 hpi [70]. (**F**) Data of qRT-PCR analysis for *BLGH_02703, BLGH_06013, BLGH_00851*, and *BLGH_06054* (up-regulated in both isolates; see Supplementary Figure 7B for additional genes) for the isolates *B. hordei* K1_AC_ (green), SK1 (maroon), SK2 (orange), and SK3 (purple) after inoculation of barley *mlo*-3. The x-axis shows the time-point after inoculation (0, 6, and 18 hpi), the y-axis displays relative transcript abundance calculated by ΔCt analysis. Data shown are based on *n* = 3 biological replicates with 3 technical replicates each.

Between 282,167 (3.5%) and 1,472,273 (9.9%) RNA-seq read pairs mapped to the manually annotated *B. hordei* reference genome DH14 v4 [27, 70] (Supplementary Table 1). The expression profiles of *B. hordei* SK1 and K1_AC_ were indistinguishable at 6 hpi but separated by fungal isolate at 18 hpi (Figure 2B). Accordingly, while we did not identify DE genes at 6 hpi, 127 genes were significantly (*P*_adj_ < 0.05; logFC > |1|) up-regulated and two genes down-regulated in *B. hordei* SK1 at 18 hpi compared to *B. hordei* K1_AC_ (Figure 2C, Supplementary Figure 6, and Supplementary Tables 5 and 6). Among the 127 up-regulated genes, 95 (74.8%) code for proteins that harbor a canonical secretion signal (mostly putative effectors; Supplementary Table 7). The two downregulated genes were *BLGH_02703* and *BLGH_06013* (see below).

Next, we compared the expression profiles of the 129 differentially expressed genes in SK1 throughout the asexual life cycle in *B. hordei* on wild-type barley (cv. Margret) plants. We took advantage of a publicly available RNA-seq dataset of *B. hordei* K1_AC_ generated at six time points of the fungal infection cycle, ranging from spore germination to sporulation, i.e., 0, 6, 18, 24, 72, and 120 hpi [70]. Of the 127 genes formally up-regulated in SK1 when grown on the *mlo*-3 mutant, 116 are usually expressed at 18 hpi in K1_AC_ on a compatible host (Figure 2D, Supplementary Figure 7A; examples are shown in Figure 2E and Supplementary Figure 7B), suggesting failure of the *mlo*-avirulent isolate K1_AC_ to induce these prior to or co-incident with haustorium establishment on barley *mlo* mutant plants. Of the remaining eleven genes formally up-regulated in SK1, six were likewise induced by K1_AC_ on the compatible host, but at later time points (“SK1 early”; Figure 2D), while another five were specifically expressed in*B. hordei* but not in K1_AC_ during the fungal infection cycle (“SK1-specific”; Figure 2D; Supplementary Figure 7B). These genes encoded putative secreted effector proteins (*BLGH_00850, BLGH_00851, BLGH_04339*, and *BLGH_07099*) and a gene of unknown function (*BLGH_05405*; Supplementary Figure 7B; Supplementary Table 7). We validated the expression profiles of eight DE genes, including four which were specifically transcribed in *B. hordei* SK1 (“SK1-specific”; Figure 2D) and the two down-regulated genes (*BLGH_02703* and *BLGH_06013*; “K1-specific”; Figure 2D), by quantitative reverse transcription polymerase chain reaction (qRT-PCR). We found that most of these genes behaved like in SK1 in the *B. hordei* strains SK2 and SK3, pointing to a common transcriptomic signature during pathogenesis in the three *mlo*-virulent isolates (Figure 2F, Supplementary Figure 8). Based on the shared profile of DE genes, we hypothesize that the five genes specifically expressed in *B. hordei* SK1, SK2, and SK3 may play a pivotal role in enabling *mlo* virulence in *B. hordei*.

### Three genes are co-affected by genomic events in the *mlo*-virulent *B. hordei* isolates

Next, we explored whether genomic alterations occur between the three *B. hordei* SK isolates and their parental meta-population (*B. hordei* K1_AC_) using whole-genome shotgun sequencing (Supplementary Table 8). We queried these genomes after at least 50 asexual generations when the *mlo* virulence phenotype had stabilized, since we expected purifying selection will have favored adaptive mutations to dominate within the fungal populations at this time. Compared to the near-chromosome level reference genome sequence of *B. hordei* DH14 [27], K1_AC_ and SK1, SK2, and SK3 shared 140,639 single nucleotide variants (SNVs), 2,137 of which were predicted to vary between the isolates. We manually confirmed 84 unambiguous polymorphisms of which 81 were intergenic and three affected genes (Figure 3A, Table 1, and Supplementary Table 9). SK2 and SK3 only differed in one intergenic SNV, implying that the two isolates are genetically near-identical. The other variations include an SNV in *BLGH_06723* (encoding a putative conserved RBR-family E3 ubiquitin ligase), causing the change of glutamine-49 to lysine in all three *mlo*-virulent isolates, and a three-base-pair (bp) deletion in gene *BLGH_06013* in part of the SK1 population, causing the in-frame loss of lysine-445 (*BLGH_06013*^ΔK445^; Figure 3B). *BLGH_06013* encodes the orthologue of *Aspergillus fumigatus* Af293 *medA* (medusa; BLASTP query cover 75%, identity 43.7%, E value 3E-112; GenBank accession XP_755658.1; Supplementary Figure 9). The above-mentioned lysine-445 (lost in BLGH_06013 in part of the SK1 population) appears to represent a positionally conserved amino acid in nearly all fungal medA orthologues analyzed (Supplementary Figure 10 and 11).

**Table 1.** Mutational events detected in *B. hordei* SK1, SK2, and SK3.

| Gene | *BLGH_06013* | *BLGH_06723* | *BLGH_02703* | *BLGH_05230*<br>*BLGH_05231*<br>*BLGH_05232* | *BLGH_05233* |
| --- | --- | --- | --- | --- | --- |
| Encoded protein | medA-like transcriptional regulator | E3 ubiquitin ligase | Unknown protein, *Blumeria*-specific | Nuclear transport factor (karyopherin) / aminopeptidase / glutathione peroxidase | Sgk2-like serine-threonine protein kinase |
| ***B. hordei* isolate** |  |  |  |  |  |
| K1<sub>AC</sub> | n.v.<sup>a</sup> | n.v. | n.v. | 2x coverage<sup>b</sup> | 3x coverage<sup>b</sup> |
| SK1 | Absent (partially) ΔK<sup>445</sup> (partially) | Q<sup>49</sup>K | Absent (partially) | 2x coverage<sup>b</sup> | 3x coverage<sup>b</sup> |
| SK2 | Absent | Q<sup>49</sup>K | Absent | 1x coverage<sup>b</sup> | 2x coverage<sup>b</sup> |
| SK3 | Absent | Q<sup>49</sup>K | Absent | 1x coverage<sup>b</sup> | 2x coverage<sup>b</sup> |

<sup>a</sup> n.v., no variation compared to the *B. hordei* reference isolate DH14 v4 [27].<sup>b</sup> relative to the *B. hordei* reference isolate DH14 v4.

**Fig. 3.**
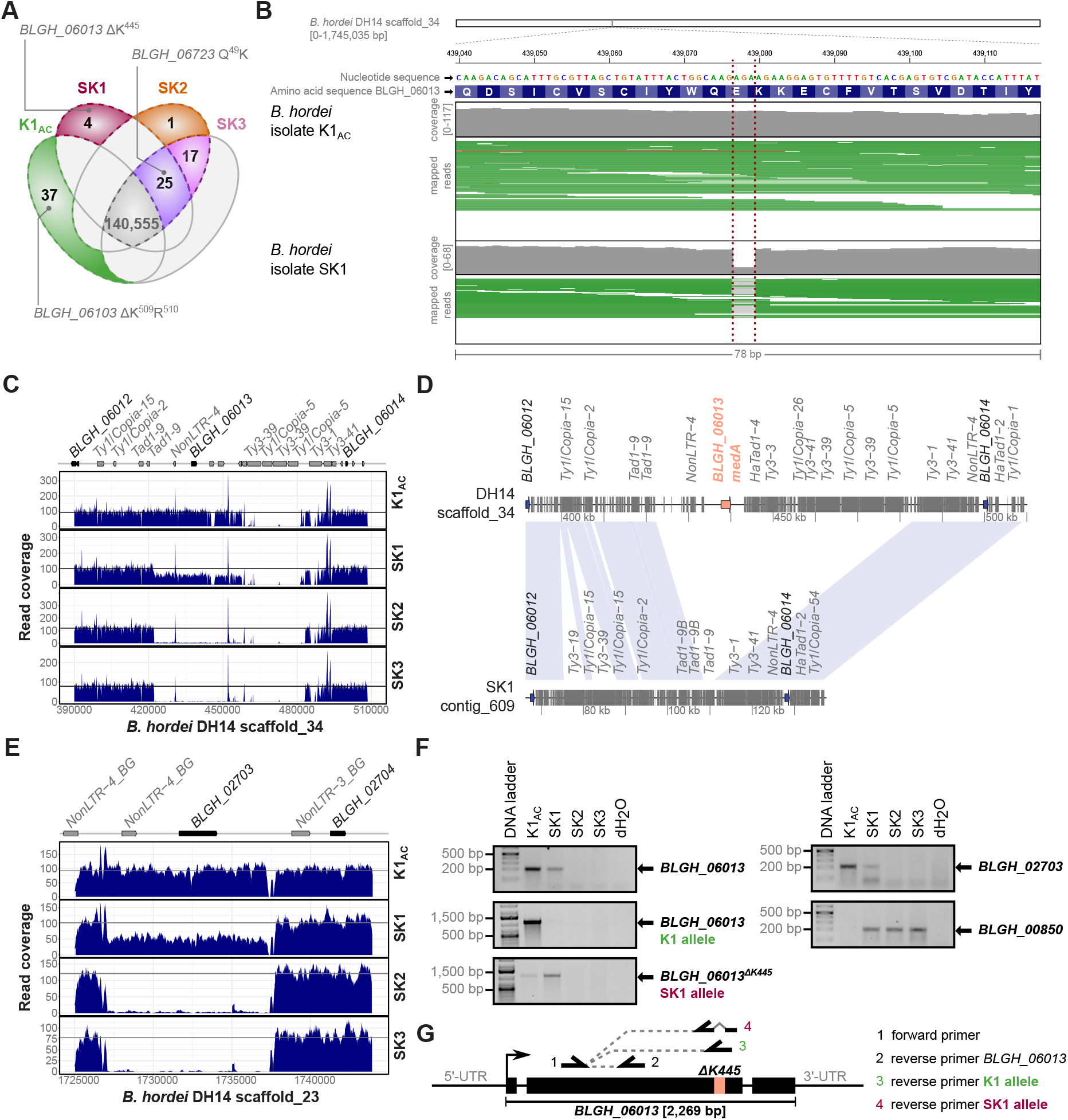
Loss of genes *BLGH_06013* and *BLGH_02703* in *mlo*-virulent *B. hordei* isolates. We performed high-throughput whole-genome DNA sequencing with the isolates *B. hordei* K1_AC_, SK1, SK2, and SK3. (**A**) Venn diagram summarizing single nucleotide variants (SNVs) occurring in *B. hordei* K1_AC_, SK1, SK2, and SK3 relative to the reference genome of *B. hordei* DH14 v4. All variants were confirmed by manual inspection, except the SNVs common in all four isolates (grey). SNVs affecting coding sequences are indicated. (**B**) SNVs were detected with Freebayes using an optimized pipeline for *Blumeria* [6] and visualized using the IGV browser [76]. The red lines highlight a variation (3-bp deletion) in *B. hordei* SK1 compared to K1_AC_. White bar, scaffold_34 of the *B. hordei* DH14 reference assembly; below, the nucleotide and amino acid sequence of *BLGH_06013* are shown. Mapping coverage is shown in grey, individual mapped reads are displayed in green, gaps in light grey. (**C**) Mapping coverage of *B. hordei* DH14 scaffold_34:390,000-510,000 of *B. hordei* K1_AC_, SK1, SK2, and SK3, including the locus of *BLGH_06013*. The x-axis shows the position on the scaffold, the y-axis read coverage, and transposable elements are indicated at the top. The black line shows the average coverage for the respective isolate, site-specific coverage is displayed in dark blue. (**D**) Local synteny plot of scaffold_34:390,000-510,000 (*BLGH_06013* locus in orange). Genes are indicated by blue arrows and transposable elements by grey blocks. *B. hordei* DH14 scaffold_34 was assembled in *B. hordei* SK1 (contig_609) using nanopore MinION sequencing. (**E**) Mapping coverage of *B. hordei* DH14 scaffold_23:1,725,000-1,750,000 of *B. hordei* K1_AC_, SK1, SK2, and SK3, including the locus of *BLGH_02703*. Displayed as in (**C**). (**F**) Genotyping PCRs for *BLGH_02703* and *BLGH_06013* alleles using genomic DNA of *B. hordei* K1_AC_, SK1, SK2, and SK3, respectively. The gene *BLGH_00850* served as a positive control for PCR amplification. DNA Ladder, 1 kb plus (Invitrogen-Thermo Fisher, Waltham, MA, USA). (**G**) Primer locations for genotyping of *BLGH_06013*. Oligonucleotides are listed in Supplementary Table 12.

We then calculated the average read coverage of all genes from the reference annotation [27], revealing two gene losses and two instances of copy number variation in the SK isolates (Supplementary Figure 12). One loss affected *BLGH_06013* and was due to deletion of an almost 40-kb region compared to *B. hordei* K1_AC_ at scaffold_34 (422,240-451,834). While this region was completely absent in *B. hordei* SK2 and SK3, SK1 exhibited about 50% loss of coverage, indicating a deletion present in part of the population (Figure 3C). We reassembled scaffold_34 of SK1 using nanopore MinION sequencing (Supplementary Table 10), which confirmed the deletion in a subset of the population with this long-read sequencing technology and revealed the apparent *de novo* insertion of a *Tad1-9* retrotransposon at this site, replacing the 40-kb region (Figure 3D). The second gene loss concerned *BLGH_02703* and was due to an 8-kb deletion in scaffold_23:1,727,078-1,735,005, where the reads again exhibited reduced (approx. 50%) coverage in SK1 and complete absence of the locus in SK2 and SK3 (Figure 3E). *BLGH_02703* has no similarity to known proteins in the NCBI database except in the closely related wheat powdery mildew pathogen, *B. graminis* f.sp. *tritici* (EPQ63962; BLASTP query cover 92%, E value 2e-154, 68.3% sequence identity and f.sp. *triticale* (CAD6506008; query cover 85%, E value 2e-166, 68.7% sequence identity), and otherwise has weak similarity to MSCRAMM family adhesin clumping factor ClfA, K^+^-dependent Na^+^/Ca2^+^ exchanger, and a guanine nucleotide exchange factor (GEF) for Rho/Rac/Cdc42-like GTPases (Supplementary Table 11). Both regions showing deletions in the SK isolates (in scaffolds 23 and 34) were flanked by long terminal repeat (LTR)-type transposable elements such as *Tad1, Ty3, Ty1/Copia*, and *Non-LTR4* (Figure 3C and 3E). Partial absence of *BLGH_02703* (*Blumeria*-specific) and *BLGH_06013* (*medA*) within the *B. hordei* SK1 population likely accounts for the reduced transcript accumulation of these two genes seen in RNA-seq analysis at 18 hpi (Figure 2D), since expression of both genes typically peaks at this time point in K1_AC_ on a compatible barley host plant (Figure 2F). We did not find the *BLGH_02703, BLGH_06013*, and ***BLGH_06723*** polymorphisms in the naturally *mlo*-virulent RACE1 isolate (Supplementary Figure 13), suggesting that its virulence is mechanistically different from *B. hordei* SK1 and SK2/3.

The copy number variations we detected in the SK isolates relative to the *B. hordei* DH14 reference genome [27] were limited to two loci (Supplementary Figure 12 and 14). Scaffold_27:312,420-348,900 encompasses *BLGH_05230, BLGH_05231*, and *BLGH_05232*, coding for a karyopherin nuclear transport factor, an aminopeptidase, and a glutathione peroxidase, respectively. The region exhibited approx. 2-fold coverage by sequence reads of isolates K1_AC_ and SK1 compared to the *B. hordei* DH14 reference genome and isolates SK2 and SK3. Likewise, sequence coverage indicated that the neighboring gene *BLGH_05233*, encoding a Sgk2-like serine-threonine kinase [48] and assembled as one copy in DH14, was represented by two copies in isolates *B. hordei* SK2 and SK3 instead of three copies in isolates *B. hordei* K1_AC_ and SK1. Scaffold_8:1,787,835-1,794,235, containing *BLGH_00850*, whose encoded protein is 36.5% similar to the protein encoded by the adjacent gene *BLGH_00851* (*CSEP0327*, an effector candidate) and thus possibly a diverged copy thereof, was recently (probably in 2018) lost in our locally propagated *B. hordei* K1_AC_ population (*B. hordei* K1_AC_) but not in the original *B. hordei* K1_AC_ population (*B. hordei* K1_CGN_) and also not in the SK descendants derived from *B. hordei* K1_AC_ (see below; Supplementary Figure 14B and 15). We confirmed the presence-absence polymorphisms of *BLGH_02703* (*Blume-ria*-specific) and *BLGH_06013* (medA) by genotyping PCR and the C^222^A nucleotide exchange in *BLGH_06723* (E3 ubiquitin ligase) by Sanger sequencing of the PCR products (Figure 3F and Supplementary Figure 16). We summarized all genomic alterations detected in the three *B. hordei* SK isolates in Table 1.

### Genomic variations emerge rapidly in the context of selection for *mlo* virulence in *B. hordei*

We conducted whole-genome re-sequencing of *B. hordei* isolates SK2 and SK3 at distinct stages of the selection process, i.e., at five, ten, and 50 asexual generations under selection pressure on barley *mlo* mutant plants (Supplementary Table 8). Most genomic variants described above were fixed in the *mlo*-virulent *B. hordei* populations and the respective parental alleles were not detected any longer at generation 50 (see above). To track the emergence and trajectory of the novel mutant alleles, we analyzed their frequency at asexual generation 5 and 10 (Supplementary Figure 17). Of the 84 confirmed genomic variations (Supplementary Table 9; see also above), 30 were already detectable at low frequencies in SK2 and 32 in SK3 at generation five, and their frequencies within the populations gradually increased to full penetrance by generation 50. By contrast, 30 variations unique to K1_AC_ but absent in SK1, SK2, and SK3 were also present at high frequencies at generations five and ten, albeit with gradually decreasing frequencies. Likewise, the haplotype of the locus containing *BLGH_00850* (Supplementary Figure 14B) was already detectable at low coverage in SK2 at generation 10 and SK3 at generation 5, and its frequency increased to full penetrance by generation 50 (Supplementary Figure 18A). Another 7 SNVs uniquely detected in K1_AC_ were either absent or only detectable at frequencies <1% at generations 5 and 10 in SK2 and SK3, indicating their counter-selection in these *mlo*-virulent isolates. By contrast, re-sequencing of K1_AC_ after five years (in 2023, equivalent to ca. 250 asexual generations) confirmed that the 37 variations uniquely found in K1_AC_ remained fixed in the *B. hordei* K1_AC_ metapopulation (Supplementary Figure 17A). Overall, the parallel gradual emergence or disappearance of alternative alleles in SK2 and SK3 suggest a sub-population that pre-existed at low frequency in K1_AC_ and that was positively selected and gave rise to the *mlo*-virulent strains.

Interestingly, neither of the variations observed for *BLGH_06013* (medA) could be observed at asexual generations 5 and 10 in SK2/SK3 (Supplementary Figure 17B). The 3-bp deletion present in SK1 could not be found, and the read coverage of *BLGH_06013* was high at both time points. Similarly, read coverage for the gene *BLGH_02703* (*Blumeria*-specific) was high as well at generation 5 and 10 (Supplementary Figure 18B), suggesting that the losses of *BLGH_06013* and *BLGH_02703* occurred after generation 10 but were nonetheless fixed in the populations by generation 50. In summary, the analysis of the SK isolates at various stages of the selection process supports the rapid initial emergence of genomic variants and a gradual shift in their allele frequencies towards the trait of stable *mlo* virulence.

### Loss of *BLGH_02703* is dispensable for *mlo* virulence of *B. hordei* SK1

We next aimed to address whether the *mlo* virulence phenotype depends on the loss of genes *BLGH_02703* (*Blumeria*-specific) and/or *BLGH_06013* (medA), which are partially absent or mutated in the *B. hordei* SK1 population and both absent in SK2 and SK3 (Figure 3, Table 1). Given the lack of reliable genetic tools for obligate biotrophic plant pathogens including powdery mildews, we took advantage of the non-homogenous SK1 population to isolate single spore-derived and PCR-validated *B. hordei* genotypes. We succeeded in separating individual colonies that either carry or lack *BLGH_02703* (*Blumeria*-specific) in combination with either the *BLGH_06013* (medA) deletion or the *BLGH_06013*^ΔK445^ variant (three different genotype combinations; Figure 4A). The entry success of these genotypes on *mlo* mutant leaves ranged from 5-12%, and there was no statistically significant difference in this respect between these isolates (Figure 4B). By contrast, reminiscent of the original SK1 isolate (Figure 1D), SK2 and SK3 as well as all SK1-derived genotypes showed reduced entry access on barley wild-type (*Mlo* genotype) leaves (Figure 4C). Transient silencing of *BLGH_02703, BLGH_06013*, and *BLGH_06723*, separately or in combination, by particle bombardment-mediated host-induced gene silencing [66] did not confer virulence to *B. hordei* K1_AC_ on barley BCI *mlo*-3 leaves (Supplementary Figure 19). Taken together, these data indicate that the loss of *BLGH_02703* (*Blumeria*-specific) is not required for *B. hordei* to acquire *mlo* virulence.

**Fig. 4.**
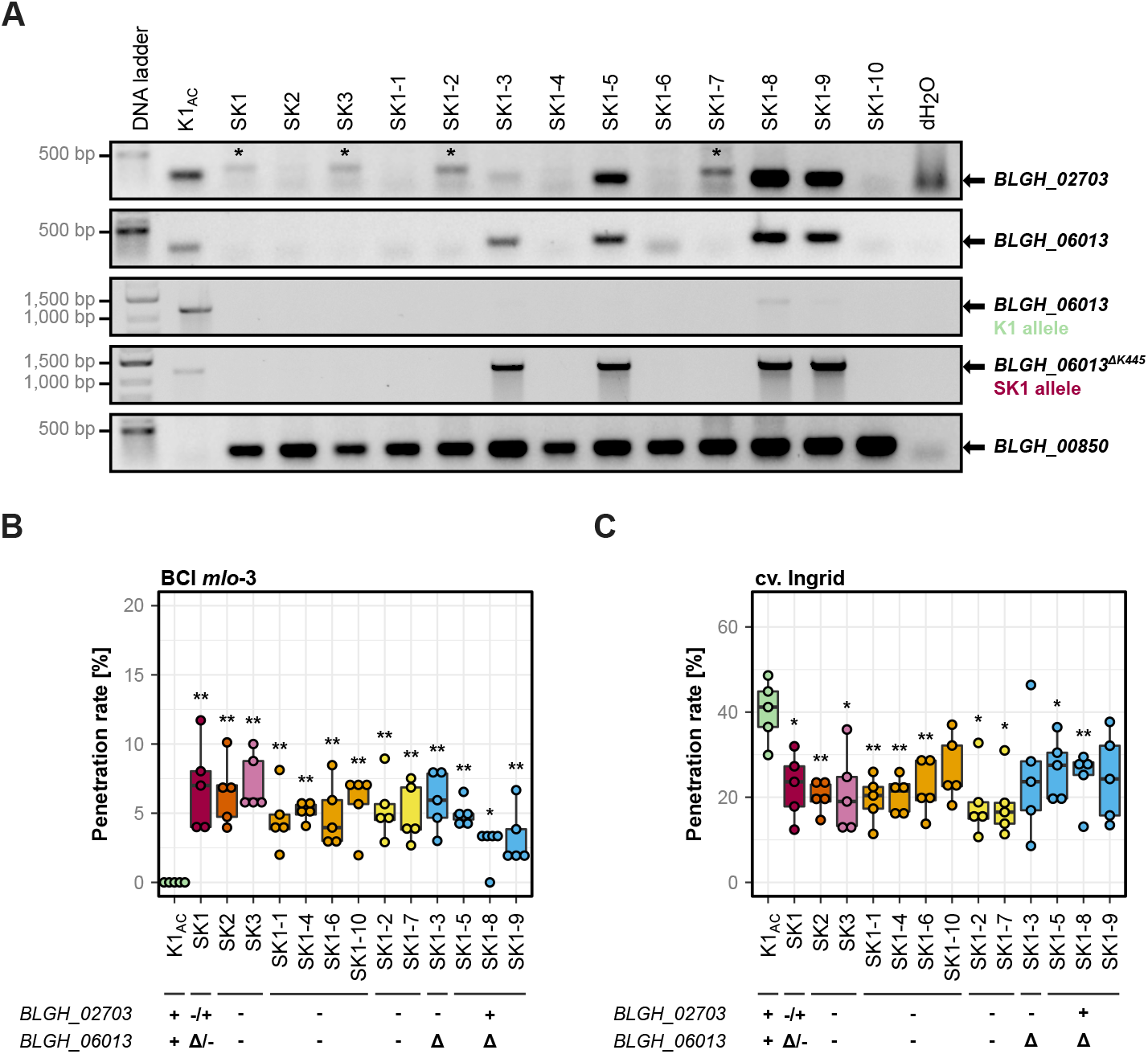
Loss of *BLGH_02703* is dispensable for *mlo* virulence in *B. hordei* SK1. Using the *B. hordei* SK1 meta-population, we captured several defined mutant genotypes and combinations thereof as single spore isolates. (**A**) Genotyping PCR for *BLGH_02703* and *BLGH_06013* with three different primer pairs as indicated in Figure 3F on genomic DNA isolated from various *B. hordei* isolates. Asterisks indicate a sporadically occurring non-specific PCR product in case of the *BLGH_02703* PCR; since the origin of these bands is unclear, lines with these bands are indicated in yellow in the boxplots in panels (**B**) and (**C**). Gene *BLGH_00850* served as a positive control for PCR amplification. (**B** and **C**) Penetration success [%] of conidia from *B. hordei* isolates K1_AC_ (green), SK1 (maroon), SK2 (orange), SK3 (purple), and the ten isolates (light orange, yellow, and blue) analyzed in (**A**) on barley BCI *mlo*-3 (**B**) and cv. Ingrid (*Mlo*) (**C**). Genotypes are color-coded as indicated below the panels (Δ indicates *BLGH_06013*^Δ^^K445^). Data are based on *n* = 5 independent replicates. At least 100 interactions per leaf and three leaves were scored for each replicate. Asterisks indicate statistically significant differences to K1_AC_ at *, *P* < 0.05 and **, *P* < 0.01, according to Kruskal-Willis and Mann-Whitney-Wilcoxon statistical tests.

### Loss of *BLGH_06013* (medA) does not affect conidia morphology in *B. hordei*

The transcription factor medA is required for the formation of normal conidia in the ascomycete *Aspergillus fumigatus*, as *A. fumigatus medA* mutants display aberrations in conidia number and shape [32, 2]. We therefore wondered whether the mutation(s) in *BLGH_06013* (*medA*) would likewise affect the morphology of asexual spores in the *B. hordei* SK isolates. We therefore wondered whether the mutation(s) in *BLGH_06013* (*medA*) would likewise affect the morphology of asexual spores in the *B. hordei* SK isolates. While we observed uniform oval-shaped conidiospores in the case of *B. hordei* K1_AC_, the *mlo*-virulent isolates SK1, SK2, and SK3 showed the presence of markedly elongated ellipse-shaped conidiospores (Figure 5A). We, therefore, assessed essential conidia size parameters (length, width, and area) of the SK isolates in comparison to *B. hordei* K1_AC_. All tested *mlo*-virulent genotypes (SK1, SK2, SK3, and the mutant combinations described above) had significantly increased conidia length (median 32-36 µm) compared to the parental isolate *B. hordei* K1_AC_ (median 23 µm; Supplementary Figure 20). The same applied, in tendency, to conidia area (approx. 213 µm^2^ vs. 160 µm^2^), while conidia width of the SK isolates was indistinguishable from *B. hordei* K1_AC_ conidia (both approx. 11 µm; Supplementary Figure 20). However, when we assessed conidia shape of the *B. hordei* K1_CGN_, A6, DH14, and RACE1, we noticed that all these isolates had conidia longer than 30 µm on average, similar to SK1, SK2, and SK3 (Figure 5B). Like *B. hordei* K1_AC_, all of these isolates carried the wild-type allele of *BLGH_06013*, suggesting that the *B. hordei medA* ortholog is not responsible for the aberrant shape of conidia in *B. hordei* K1_AC_, and that conidia morphology is not correlated with *mlo* virulence in isolates SK1, SK2, and SK3. The discrepancy in spore morphology between *B. hordei* K1_AC_ and all other tested *B. hordei* isolates points towards a spontaneous mutation in K1_AC_, affecting this trait that must have occurred and became fixed in the population after the isolation of the SK isolates.

### *mlo* virulence may cause an overall fitness penalty in *B. hordei*

We had noticed early on that the aberrant virulent isolate *B. hordei* SK1 displayed a reduced entry rate compared to K1_AC_ on barley cultivars carrying a functional *Mlo* gene (Figure 1D). To test the viability of conidia, we allowed the conidia of *B. hordei* K1_AC_, SK1, SK2, SK3, and the various genotypes isolated from the SK1 meta-population to germinate on agar medium. Under these *in vitro* conditions, around 50-75% of conidia formed germ tubes. The majority of the SK isolates displayed normal or slightly enhanced germination rates (approx. 50-70%; Figure 5C), in-dicating that germination and formation of the primary germ tube remained largely unaffected between *mlo*-virulent isolates and by the aberrant conidia shape in K1_AC_. Then, we analyzed a set of barley cultivars susceptible to *B. hordei* K1_AC_ and carrying a functional *Mlo* allele to compare the entry success of the two isolates *B. hordei* K1_AC_ and SK1. Intriguingly and consistent with our other experimental data (Figure 1D and Figure 4C), *B. hordei* SK1 showed a significantly reduced entry success with a decrease of about 5-15% compared to *B. hordei* K1_AC_ on these cultivars, suggesting a fitness penalty for *B. hordei* SK1 (Figure 5D). To assess whether *B. hordei* K1_AC_ would rapidly outcompete *B. hordei* SK1 as a consequence of this fitness penalty, we performed competition experiments. However, *B. hordei* K1_AC_ did not outcompete *B. hordei* SK1 in these settings, and *mlo*-virulent *B. hordei* isolates did not lose their *mlo* virulence after 12 generations of propagation on wild-type barley leaves (cv. Ingrid; *Mlo* genotype), i.e., in the absence of the selective pressure (Supplementary Figure 21).

**Fig. 5.**
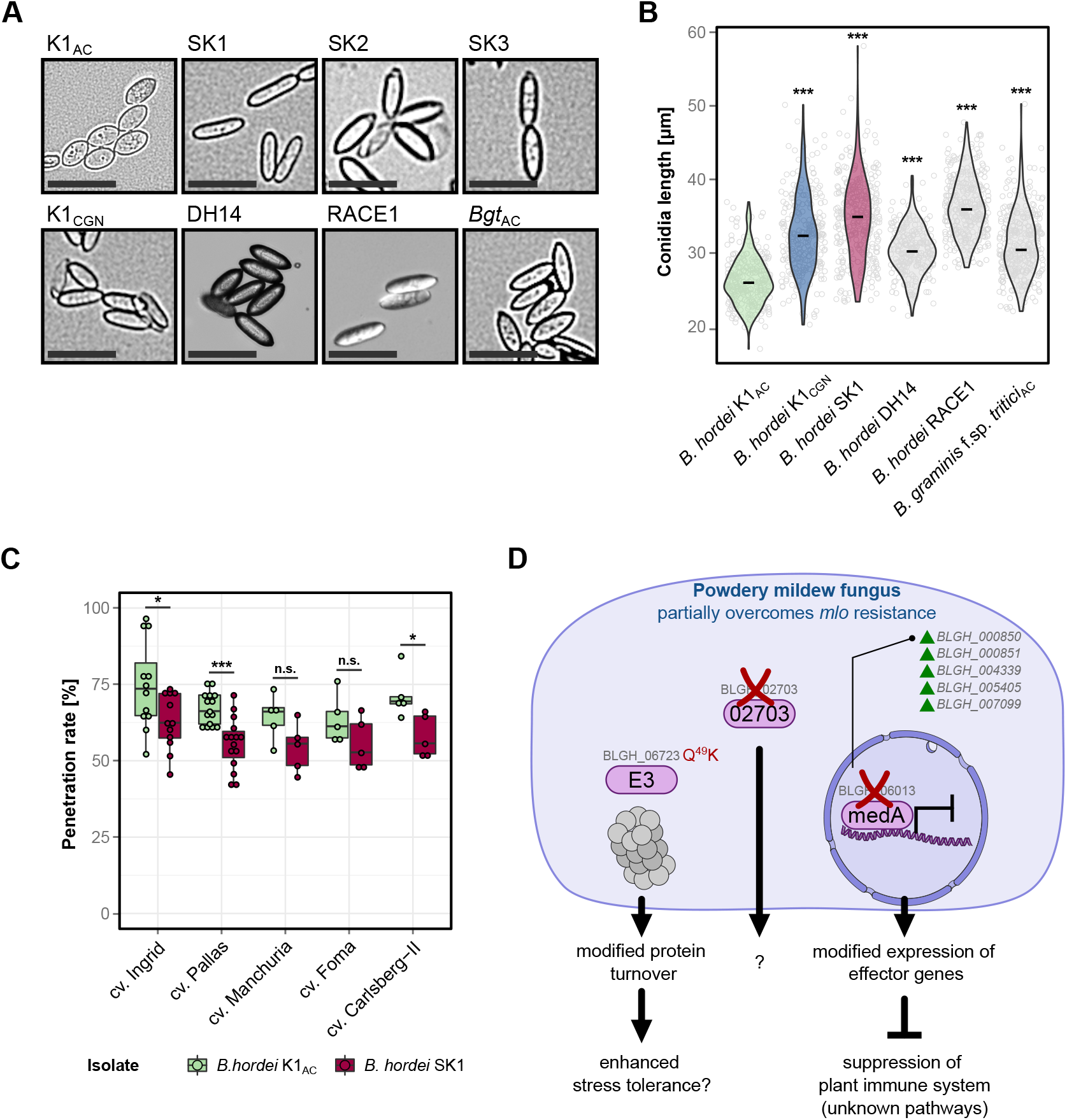
The loss of *BLGH_06013* does not affect conidiospore morphology. (**A**) Representative micrographs of conidia (brightfield); scale bar: 50 µm. Conidia were obtained from *B. hordei* K1_AC_, SK1, SK2, SK3, K1_CGN_, DH14 (courtesy by Pietro Spanu, Imperial College London), RACE1, and *B. graminis* f.sp. (**B**) *tritici*. The violin plot shows the length of conidia [µm] for *B. hordei* K1_AC_, SK1, K1_CGN_, DH14, RACE1, and *B. graminis* f.sp. *tritici* isolate Aachen (*Bg tritici* _AC_). At least 300 conidia were measured for each isolate. A statistical test according to Mann-Whitney-Wilcoxon was performed to estimate differences to K1_AC_; ***, *P* < 0.001. (**C**) Penetration success [%] of conidia from *B. hordei* K1_AC_ (green) and SK1 (maroon) on barley cultivars (*Mlo* genotype) Ingrid (n = 12), Pallas (*n* = 15), Manchuria, Foma, and Carlsberg-II (*n* = 5 independent replicates each). Statistical analysis was performed comparing SK1 with K1_AC_ on the respective cultivar using Kruskal-Willis and Mann-Whitney-Wilcoxon statistical tests; *, *P* < 0.05; **, *P* < 0.01; ***, *P* < 0.001; n.s., not significant. (**D**) Simplified model for partial *mlo* virulence in the barley powdery mildew pathogen. Multiple pathways contribute to *mlo*-based resistance against powdery mildew pathogens. Our experiments demonstrated that *mlo* virulence in *B. hordei* SK1 is additive to suppression of resistance by loss of known defense pathways, suggesting a different mechanism that leads to successful infection. We identified three proteins to be associated with the *mlo* virulence in *B. hordei* SK1 and SK2/3: the E3 ubiquitin ligase BLGH_06723, which may be involved in protein turnover, the transcriptional regulator BLGH_06013 (medA), whose absence or non-functionality may allow the activation of several genes including candidate effector-encoding genes, and a third *Blumeria*-specific protein of unknown function (BLGH_02703). We postulate that these three proteins alter the infection program of *B. hordei* SK1 and SK2/3 to collectively enable infection of *mlo*-resistant barley. Purple ovals symbolize the three proteins found to be lacking/mutated in the three *B. hordei* SK1 isolates. The schematic was generated using icons from https://bioicons.com: proteasome icon by jaiganesh (https://github.com/jaiganeshjg); nucleus-full-3d and dna-5 icon icons by Servier (https://smart.servier.com/).

## Discussion

Experimental evolution can be a powerful tool to study genome evolution in plant-associated fungal microbes [60]. Using this approach, we discovered a single amino acid exchange in one gene and the loss of two genes co-occurring in all three *B. hordei* SK isolates (Table 1). This mutational spectrum correlates with the gain of virulence on otherwise highly powdery mildew-resistant barley *mlo* mutant plants, which confer a type of broad-spectrum resistance that mechanistically differs from isolate-specific immunity conferred by NLR proteins [39, 69]. Different from mutations leading to the loss of NLR-mediated isolate-specific resistance, none of these genes code for effectors but rather a RBR-family E3 ubiquitin ligase (*BLGH_06723*), a medA-like transcriptional regulator (*BLGH_06013*), and a *Blumeria*-specific protein of unknown function (*BLGH_02703*). The Q^49^K sub-stitution in *BLGH_06723* does not affect a conserved amino acid and is not located in an annotated functional domain of the protein (Supplementary Figure 22). We can, however, not exclude that this missense mutation leads to either a non-functional version or a gain-of-function variant, or, alternatively, affects the stability and accumulation levels of the protein. Our analyses based on the segregating SK1 population demonstrated that the lack of *BLGH_02703* is dispensable for both *mlo* virulence and the conidiospore morphology phenotype (Figure 4 and 5). Likewise, the copy number variation of genes *BLGH_05230, BLGH_05231, BLGH_05232*, and *BLGH_05233* is restricted to *B. hordei* SK2 and SK3 (Table 1). These variations are thus unlikely to be causative for *mlo* virulence. This leaves the Q^49^K substitution in *BLGH_06723* and/or the lack of *BLGH_06013* as the most probable mutations conferring partial *mlo* virulence in the *B. hordei* SK isolates. The lack of genetic tools for the obligate biotrophic powdery mildew pathogens at present prevents a more rigorous testing of the candidate genes, e.g., by targeted gene knock-outs or complementation analysis.

It is intriguing that all three *B. hordei* SK isolates share an almost identical set of adaptive mutations, including the joint occurrence of a single amino acid substitution in *BLGH_06723* (Table 1). Similarly, previously isolated *B. hordei* isolates depended on three unidentified genes that unequally contributed to *mlo* virulence [3, 33]. These earlier experiments resulted in strains of three distinctive levels of virulence, suggesting at least three major adaptive mutations [82]. This observation prompts the question whether the SK isolates represent independent mutational events, and whether the detected sequence variants might pre-exist as balanced polymorphisms within the *B. hordei* K1_AC_ population or whether they are independently and convergently acquired *de novo* events that were selected for during experimental evolution. Since isolate SK1 on the one hand and SK2/SK3 on the other hand were retrieved more than two years apart from each other, and differ in the mutational spectrum detected for *BLGH_06013* (medA) and the copy number variation of *BLGH_05230, BLGH_05231, BLGH_05232*, and *BLGH_05233* (Table 1), we can assume at least two independent and convergent sets of adaptive mutational events.

By contrast, isolates SK2 and SK3 are near-identical and according to our analysis only differ by one validated SNV in an intergenic region, suggesting that these two isolates might have the same origin. *B. hordei* isolates are non-homogenous and have a complex population structure, with many different haplotypes present within a supposedly clonal isolate [6]. It may thus well be possible that a founder event for *mlo* virulence (e.g., the Q^49^K substitution in BLGH_06723) is present at low levels as a balanced polymorphism within the parental *B. hordei* K1_AC_ population, which may give rise to sporadic colony formation on *mlo* plants. One or more additional events, such as the loss of functional *BLGH_06013* (medA) might be required to stabilize virulence on *mlo* mutant plants. While we found no evidence for the BLGH_06723 Q^49^K variant to be present in the K1_AC_ population, we detected >70 SNVs including this one at asexual generation 5 after start of the experimental selection at low frequencies in both SK2 and SK3, while variants in *BLGH_06013* and *BLGH_02703* were not detectable, supporting the hypothesis of a founder population in which *mlo* virulence is later stabilized by additional events. This notion is further strengthened by the earlier experimental evolution approaches conducted by Erik Schwarzbach, who likewise observed discrete step-wise increases in *mlo* virulence in the course of the experiment [82]. Gene loss is often associated with adaptation to new host environments and host jumps in fungal pathogens [84]. Various processes could facilitate the rapid gain and loss of genes in *B. hordei*. Genome recombination due to mating (sexual reproduction) can promote genetic diversity, which for example gave rise to the emergence of *B. graminis* f.sp. *triticale* by hybridization of wheat and rye powdery mildews [62]. However, we isolated the *mlo*-virulent strains from asexually-reproducing populations. Thus, the extensive repertoire of transposable elements [27] is the most likely driver of rapid genomic changes in the fungus. We noted that in the case of *BLGH_06013* a *Tad1-9* transposon appears to have replaced the locus containing this gene in SK1 (Figure 3D), suggesting that transposition of a *Tad1* element caused genome recombination at this site, consistent with the copy-paste mechanism of retroelements [92]. However, the transposition itself would not explain the loss of a large genomic segment. While non-homologous end joining (NHEJ) is the dominant mechanism to repair double strand breaks in genomic DNA in haploid genomes [78, 14], it accounts predominantly for indels of <20 bp. Microhomologies however can cause long-distance template switching due to replication fork collapse during cell division, which can frequently occur in repetitive and AT-rich regions [9, 90]. The resulting DNA end intermediates can be stabilized and repaired by microhomology-mediated end joining (MMEJ), which in this case uses microhomologies from non-homologous templates and is therefore error-prone [13, 37, 86]. Microhomology-driven replication-based DNA repair mechanisms such as MMEJ cause large-scale insertions, deletions, and copy number variation [21, 67, 79]. Since powdery mildew genomes are enriched with repetitive elements including retrotransposons that exhibit little sequence divergence [27, 93], microhomologies likely occur frequently and could give rise to extensive structural variation in the fungus. Typically, virulence genes encoding effectors, carbohydrate-processing enzymes (CAZymes), and toxins are often found in the vicinity of transposable elements, frequently embedded within transposon-rich genome compartments [20, 4, 25]. For example, population-wide screening of the gene content in the Septoria leaf blotch pathogen *Zymoseptoria tritici* identified 599 gene gains and 1,024 gene losses, which mainly occurred in subtelomeric regions and in proximity to transposable elements. The majority of these genes encode virulence factors, secreted proteins, and enzymes involved in the biosynthesis of secondary metabolites [36]. Inaccurate microhomology-based repair of double strand breaks occurring in the transposon-rich and repetitive regions may be one major driver of copy number variation of effectors and other virulence factors.

Why do the mutational events observed in the *B. hordei* SK isolates not occur naturally in barley powdery mildew populations in the field? This might be explained by the fact that the affected isolates show reduced infection success on susceptible (*Mlo* wild-type) barley genotypes (Figure 5D). Due to the adverse effects of this adaptation, such strains may not emerge in a non-selective environment where susceptible barley genotypes are available as hosts. It is conceivable that the rotation of *mlo*-resistant and non-resistant spring and winter varieties, respectively, as currently practiced by farmers in European agriculture [41], results in the absence of constant selection pressure, thereby preventing the occurrence of natural *mlo*-virulent strains so far. Given the rapidity with which *mlo* virulence appeared under our laboratory conditions, we caution against the permanent deployment of barley *mlo* mutants without rotation to prevent the appearance of *mlo*-virulent barley powdery mildew in agricultural settings. In summary, we established experimental evolution as a novel addition to the genetic toolbox available to study obligate biotrophic pathogens such as powdery mildew fungi. This approach complements the recently established mutagenesis pipelines [6, 7]. The complex (meta-)population structures of fungal isolates [6], which confounds genomic analyses, and the lack of effective transformation systems for the validation of candidate genes nonetheless remain challenges for the work with these organisms.

## Methods

### Plant growth conditions

All plants were cultivated in SoMi513 soil (HAWITA, Vechta, Germany). Healthy barley and wheat plants were grown under a long day cycle (16 h light period at 23 °C, 8 h darkness at 20 °C) with 60-65% relative humidity (RH) at a light intensity of 105-120 µmol s^-1^ m^-2^. *Arabidopsis thaliana* plants were cultivated under a short-day cycle (8 h light period at 22 °C, 16 h darkness at 20 °C), at 80-90% RH, and a light intensity of 100 µmol s^-1^ m^-2^. For powdery mildew infection assays, the plants were transferred to isolate-specific infection chambers with a long day cycle (12 h light at 20 °C and 12 h dark period at 19 °C), ca. 60% relative humidity and 100 µmol s^-1^ m^-2^.

### Powdery mildew infection assays

One-week-old barley and wheat plants and four to five-week-old *Arabidopsis* plants (rosette size of 2-2.5 cm) were used for powdery mildew infection assays. The powdery mildew conidiospores were blown onto the plants in an infection tower; spores were allowed to settle for 10-15 min. Inoculated plants were incubated in the respective *B. hordei* infection chamber. The samples for penetration assays were bleached in 80% ethanol at 48 hpi. The leaves were submerged twice in Coomassie staining solution (45% v/v MeOH, 10% v/v acetic acid, 0.05% w/v Coomassie blue R 250; Carl Roth, Karlsruhe, Germany) for 15-20 s, and then mounted on a glass slide with 50% glycerol. The samples were evaluated by bright field microscopy. Leaves from four to five plants/genotype were scored for penetration success with 100-200 interaction sites per leaf. Penetration success is expressed as the percentage of spores forming secondary hyphae upon interaction over spores forming an appressorium only.

For the *B. hordei*-*Arabidopsis* interaction assays, the leaves were stored in Aniline blue staining solution (150 mM K_2_HPO_4_, 0.01% w/v Aniline blue; Sigma-Aldrich, Munich, Germany) in the dark overnight to stain callose, before Coomassie staining. These samples were analyzed by fluorescence microscopy with illumination by an ultraviolet (UV) lamp (bandpass 327-427 nm) and an emission filter for Aniline blue/DAPI at 417-477 nm.

### Analysis of conidia shape

Conidia were collected from the surface of susceptible barley leaves at 7 dpi using transparent Scotch^TM^ tape, which was then mounted onto a glass slide with 20 µL of tap water. Brightfield photographs were taken with the Keyence Biorevo BZ-9000 and BZII Viewer software (Keyence, Osaka, Japan) using the 10x magnification objective. Conidia shape (length and width, area, and perimeter) was determined using the ImageJ (https://imagej.nih.gov) function Analyze Particles.

### Conidia germination assay

Conidia from 7-10-days-old powdery mildew colonies were blown onto 1% agar-agar Kobe I (Carl Roth, Karlsruhe, Germany). Conidia germination was assessed via brightfield microscopy at 6 hpi, scoring the percentage of spores that formed primary germ tubes and counting the number of germ tubes on germinated conidia.

### Whole transcriptome shotgun sequencing analysis

Epiphytic fungal material was collected as described in [53] at 6 hpi (appressorium formation) and 18 hpi (early penetration). Whole transcriptome shotgun sequencing (RNA sequencing) was done by the service provider CeGaT (CeGaT, Tübingen, Germany), yielding 100-bp paired-end reads. Raw reads were trimmed using Trimmomatic v0.39 [8] and quality control of the reads was done with FastQC v0.12.1 (Babraham Bioinformatics, Cambridge, UK). HISAT2 [43] with `-max-intronlen 1000 -k 1` mapped the reads to the *B. hordei* DH14 reference genome [27] and the *H. vulgare* cv. Morex reference genome version Morex3 [61]. The SAM/BAM files were parsed with SAMtools v1.18 [52] and BEDtools v2.31.0 [71]; read counts were determined using featureCounts v2.0.1 [54] with the gene annotations for *B. hordei* DH14 v4.3 [70] and Morex3 [61], respectively. Non-expressed genes were removed with a cut-off of TPM < 1 in any sample. Differential expression analysis was performed via the limma-VOOM pipeline with cut-offs log-fold-change > |1| and *P*_adj_ < 0.05 using limma v3.56.2 [51], DESeq2 v1.40.2 [57], and EdgeR v3.56.2 [77]. Genes differentially expressed in *B. hordei* SK1 were compared with their expression pattern in the avirulent isolate *B. hordei* K1_AC_ throughout the infection life cycle using RNA-seq time-course expression data obtained previously [70]. Differential expression of selected genes was verified by qRT-PCR, performed as in [68]; primers are listed in Supplementary Table 12.

### Quantitative real-time polymerase chain reaction (qRT-PCR)

Detached barley leaves were placed on 1% agar-agar Kobe I (Carl Roth, Karlsruhe, Germany) plates containing 85 µM benzimidazole, and inoculated with *B. hordei* isolates K1_AC_, SK1, SK2, and SK3, respectively. Epidermal peelings were collected at 0, 6, and 18 hpi in three biological replicates, and flash-frozen in liquid nitrogen. RNA extraction was performed using the TRIzol protocol (Invitrogen-Thermo Fisher, Waltham, MA, USA). RNA concentration was determined using Nanodrop 2000c (Thermo Fisher Scientific) and RNA integrity was assessed on a 2% agarose gel. Genomic DNA removal was done via DNase I (RNase-free, Thermo Fisher Scientific). Complementary DNA (cDNA) was synthesized using the High-Capacity RNA-to-cDNA Kit (Applied Biosystems-Thermo Fisher) and stored at -20 °C until further use. qRT-PCR was performed using the Takyon No ROX SYBR MasterMix blue dTTP Kit (Eurogentec, Seraing, Belgium) and the LightCycler 480 II (Roche, Rotkreuz, Switzerland). 1:10-diluted cDNA was used as a template for qRT-PCR reactions. The PCR efficiency of all primers used in this study (Supplementary Table 11) was between 1.8 and 2.0 and the annealing temperature was set to 58 °C. Evaluation of expression levels of target genes in relation to the housekeeping gene *B. hordei GAPDH* [68] was performed using the ΔCT method [56], calculated as 2^-(CT(*target*) – CT(*GAPDH*))^. Each biological replicate was measured in technical triplicates.

### Genome sequencing and analysis

High molecular weight genomic DNA from barley powdery mildew conidia was generated according to [23] with the modifications indicated in [27]. DNA shotgun sequencing was performed using Illumina NovaSeq technology with 1 µg input DNA at the service provider CeGaT (CeGaT, Tübingen, Germany), yielding 150-bp paired-end reads. Longread sequencing by MinION (Oxford Nanopore Technologies, Oxford, US) technology and genome assembly of *B. hordei* SK1 were done as in [27]. Short-read sequencing data from this work and from *B. hordei* isolates sequenced previously, i.e., K1_CGN_ and A6 [35], and DH14 and long-read sequencing data from *B. hordei* RACE1 [27], were re-mapped to the *B. hordei* DH14 reference genome [27] using the function `bwa mem` of BWA v0.7.17-r1188 (Li and Durbin, 2009). Single Nucleotide Variant (SNVs), insertions, and deletions (indels) were detected with FreeBayes v1.3.7-dirty [30] and raw SNVs and indels were filtered using VCFtools v0.1.16 [17] and bcftools v1.17 (https://samtools.github.io/bcftools/bcftools.html) according to [6]. We used SnpEff v4.3t (build 2017-11-24 10:18) [15] to identify SNVs and indels in genic loci with predicted effects, and manually inspected candidate polymorphisms with Integrative Genomics Viewer (IGV) browser v2.16.1 [76]. Genome mapping coverage was determined with BEDtools v2.31.0 [71]. Synteny analysis was performed using the function `nucmer` from MUMmer v3.23 [46] and visualized using the R package genoPlotR v0.8.11 [34]. Gene losses and gene alleles were verified by *Taq* polymerase-based polymerase chain reaction (PCR) on genomic DNA; primers are listed in Supplementary Table 12.

### Phylogenetic and functional analysis

Orthologues of *Aspergillus fumigatus* and *B. hordei medA* were identified using BlastP (https://blast.ncbi.nlm.nih.gov/Blast.cgi) at E value < 1E-25. Protein alignments were done with ClustalW in MEGAX [45] and visualized using Jalview v2.11.3.2 [91]. The phylogenetic tree building was facilitated by Phylogeny Analysis at http://www.phylogeny.fr, with 100 bootstrap replications and otherwise default parameters. Functional predictions were done with Inter-ProScan (https://www.ebi.ac.uk/interpro/), NCBI CDART (www.ncbi.nlm.nih.gov/Structure/lexington/lexington.cgi), PROSITE (https://www.expasy.org/resources/prosite), and protein disorder by IUPRED3 [22]. Protein structures were visualized with YASARA (http://www.yasara.org); structural comparison of BLGH_06723 was performed against E3 ubiquitin-protein ligase parkin of *Rattus norvegicus* (10.2210/pdb4K95/pdb/ [89]).

### Statistical analysis

The statistics program R v4.3.1 [73] (R foundation, www.r-project.org) was used for data analysis, statistics, and plotting. Data analysis was supported by the packages tidyverse v2.0.0, dplyr v1.1.2, reshape2 v 1.4.4, and scales v1.2.1. Principal component eigenvalues were calculated with the function `prcomp`. Sample distances for non-metric dimensional scaling (NMDS) analysis were determined with the package vegan v2.6-4; hierarchical clustering was done with functions `dist()` and `hclust()`.

For statistical analysis, data were first assessed for normal distribution by performing visual inspection via density and Q-Q plots and normality testing with the Shapiro-Wilk method `shapiro.test()` in R. The type of data collected here was ratio data and the groups were unpaired and normality testing indicated non-normal distribution of data in all cases. Hence, we performed Kruskal-Wallis tests via `kruskal.test()` and pairwise comparison via Mann-Whitney-Wilcoxon testing using the function `pairwise.wilcox.test()` to determine statistical differences between groups. Where appropriate, we used the multcompView package function `multcompLetters()` to assign multivariate comparisons into statistical groups de-noted by letters.

Heat maps were generated with the R package gplots v3.1.3 using the function `heatmap.2`, dot plots, boxplots and violin plots were done using the R package ggplot2 v3.4.2 [94]. Venn diagrams were plotted with the online tool jvenn at https://jvenn.toulouse.inrae.fr/app/index.html [5].

## Abbreviations

BCI: backcross Ingrid
bp: base pair
DE: differentially expressed
dpi: days post-inoculation
hpi: hours post-inoculation
indel: insertion and deletion
*Mlo*: *Mildew resistance locus O*
PCA: principal component analysis
PCR: polymerase chain reaction
RNA-seq: RNA-sequencing
qRT-PCR: quantitative reverse transcriptase-polymerase chain reaction

## Data availability

All raw RNA and DNA sequencing data generated in this study are deposited at https://www.ebi.ac.uk/ena under project IDs PRJEB36770 (*B. hordei* K1_AC_ [27]) and at https://www.ncbi.nlm.nih.gov/sra under BioProject ID PRJNA639160. The draft genome assembly for *B. hordei* SK1 has been deposited at DDBJ/ENA/GenBank under the accession JAJOCF000000000.

## Code availability

We applied default parameters of the computational tools used in this study, unless indicated otherwise. We report all tools and versions.

## Availability of biological materials

Barley powdery mildew is an obligate biotrophic pathogen and cannot be cultivated *in vitro*, nor stored long-term, e.g., by glycerol stocks. Depositing the newly obtained strains in a stock center is not possible for this reason. Therefore, we keep the unique strains *B. hordei* SK1, SK2, and SK3 alive by weekly reinoculation of living barley plants. Barley leaves infected with these strains can be obtained upon request.

## Acknowledgements

Seeds of the wheat cultivars used in this study were kindly provided by the late Patrick Schweizer (IPK Gatersleben, Germany) and Beat Keller (Zurich University, Switzerland). We thank Bernd Denecke (IZKF Aachen) for permitting access to the Illumina NextSeq sequencer and the Agilent Bioanalyzer platform. We acknowledge Alexander Vogel, Bianca Reiß, and Björn Usadel (RWTH Aachen University) for access to DNA quantification via Qubit. The analysis was performed with computing resources granted by RWTH Aachen University under project ID rwth0146.

This study was supported by the RWTH Aachen Excellence Initiative [RWTH fellow grant to R.P.] and funded by the Deutsche Forschungsgemeinschaft (DFG, German Research Foundation) project number 274444799 [grants 861/14-1 and 861/14-2 awarded to R.P.] in the context of the DFG-funded priority program SPP1819 “Rapid evolutionary adaptation – potential and constraints”.

## AUTHOR CONTRIBUTIONS

R.P. and S.K. designed the study; R.P., L.F., and S.K. were responsible for experiment conception. L.P. generated *B. hordei* SK1, L.F. and M.B. *B. hordei* SK2 and SK3. S.K., B.D.L., and K.D.W. performed the pathogen assays. L.F. generated the samples for RNA-seq, S.K. analyzed the quantitative data and the RNA-seq data. S.K., L.F. and M.B. prepared high molecular weight genomic DNA of the *B. hordei* strains. L.F., M.B., and S.K. performed genome assemblies, comparative genomics, and subsequent data analysis. F.K. sampled and isolated RNA for qRT-PCR, performed qRT-PCR, and cloned the *BLGH_06013* alleles. S.K. wrote the first draft of the manuscript and S.K. and R.P. edited the manuscript; L.F. provided critical feed-back on the drafts. All authors read the manuscript and approved the final version.

## Supplementary data

**Supplementary Table 1**. RNA-seq mapping statistics.

**Supplementary Table 2**. Differential expression analysis results of *H. vulgare* BCI *mlo-3* at 6 hpi with *B. hordei* SK1 and K1_AC_.

**Supplementary Table 3**. Differential expression analysis results of *H. vulgare* BCI *mlo-3* at 18 hpi with *B. hordei* SK1 and K1_AC_.

**Supplementary Table 4**. Annotations of differentially expressed genes in *H. vulgare* BCI *mlo-3* at 18 hpi.

**Supplementary Table 5**. Differential expression analysis results of *B. hordei* SK1 compared to *B. hordei* K1_AC_ on *H. vulgare* BCI *mlo-3* at 6 hpi.

**Supplementary Table 6**. Differential expression analysis results of *B. hordei* SK1 compared to *B. hordei* K1_AC_ on *H. vulgare* BCI *mlo-3* at 18 hpi.

**Supplementary Table 7**. Annotations of differentially expressed genes in *B. hordei* SK1 at 18 hpi.

**Supplementary Table 8**. Whole genome shotgun DNA sequencing mapping statistics.

**Supplementary Table 9**. List of SNVs detected and manually inspected in K1_Aachen_, SK1, SK2, and SK3 and their allele frequencies.

**Supplementary Table 10**. Genome assembly statistics for *B. hordei* SK1 compared to publicly available *B. hordei* genome assemblies.

**Supplementary Table 11**. NCBI CDART results for the protein *BLGH_02703*.

**Supplementary Table 12**. List of oligonucleotides used in this study.

