## Supplementary figures and images for "A fungal plant pathogen overcomes *mlo*-mediated broad-spectrum disease resistance by rapid gene loss"

### Supplementary Figure 1

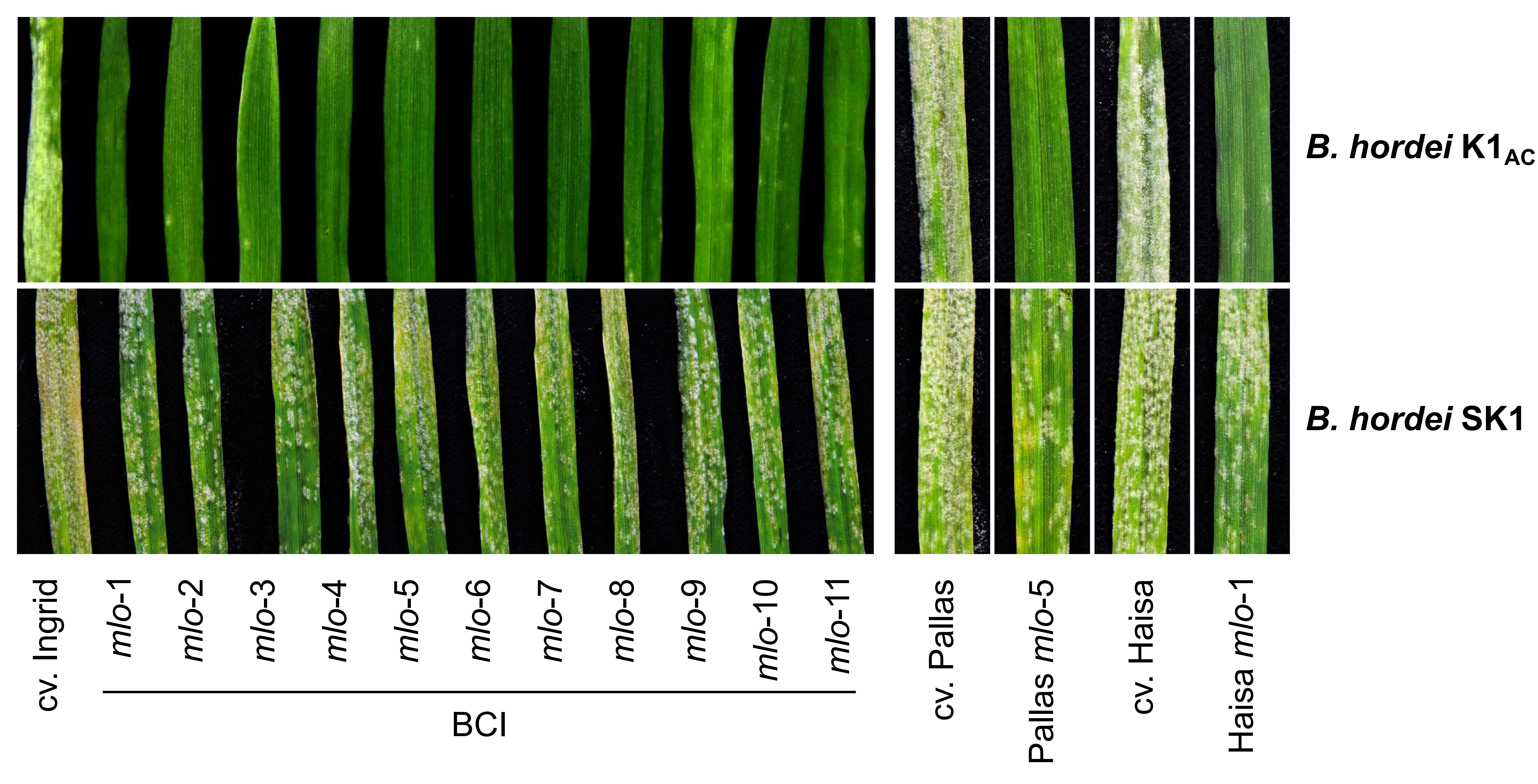

### Supplementary Figure 2

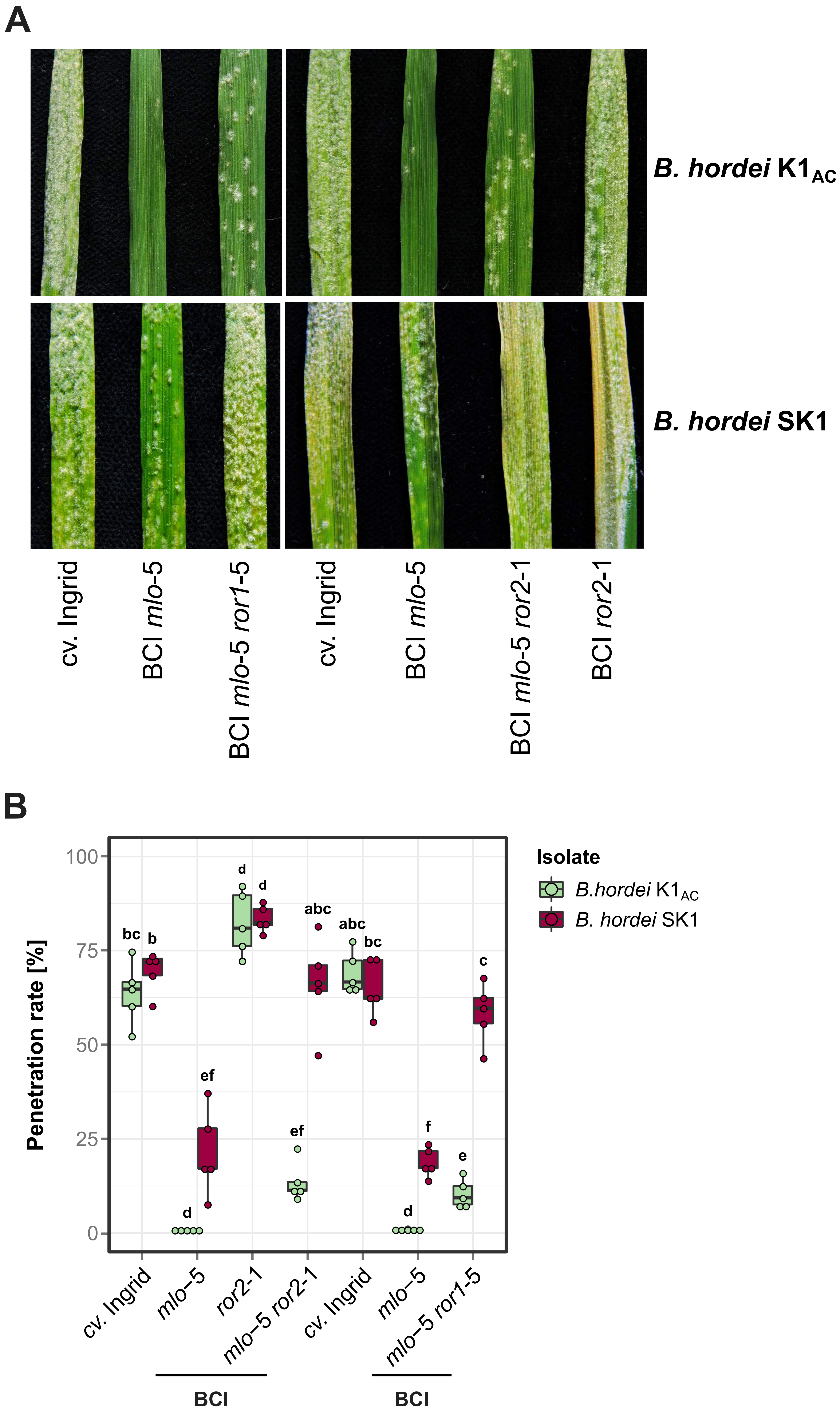

### Supplementary Figure 3

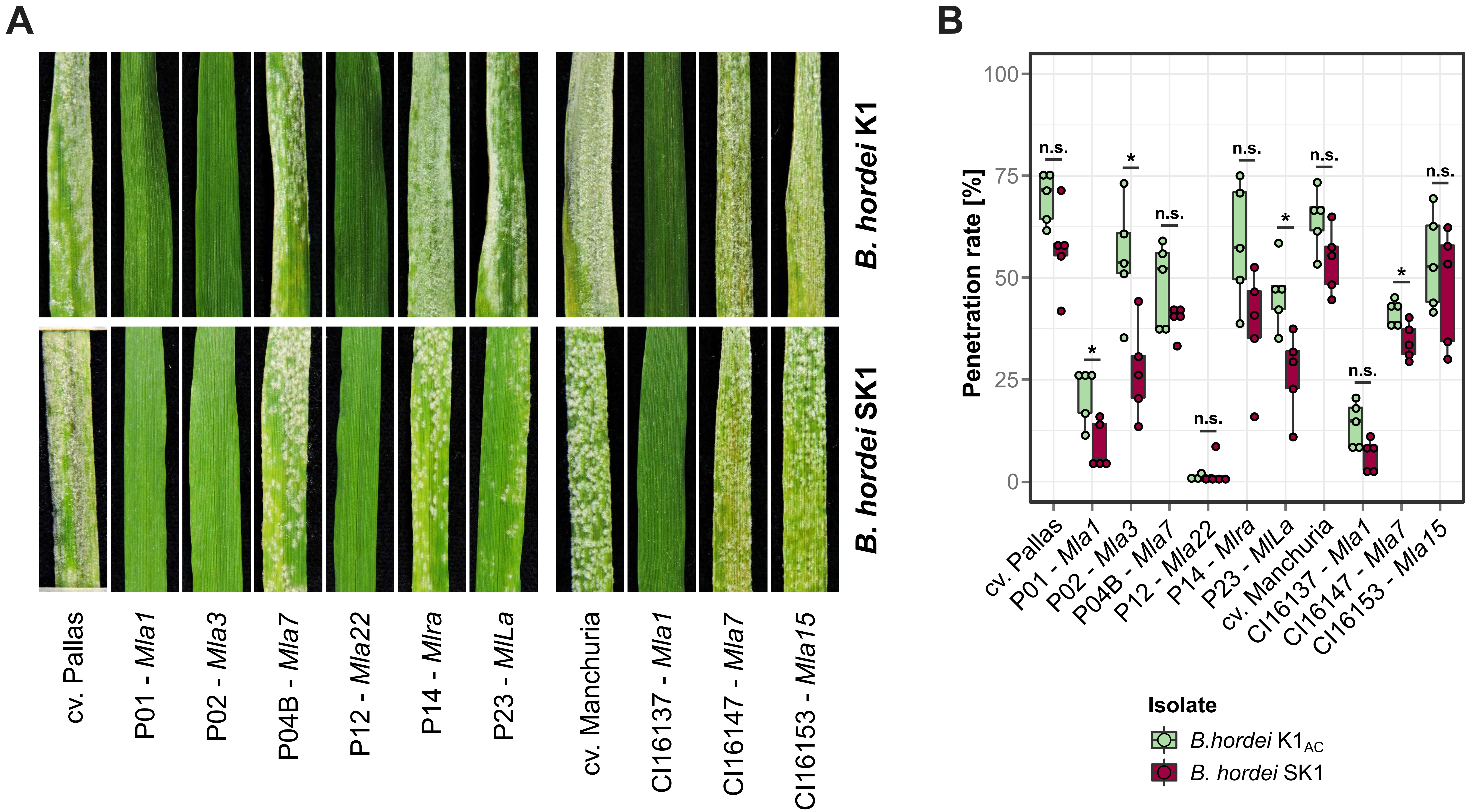

### Supplementary Figure 4

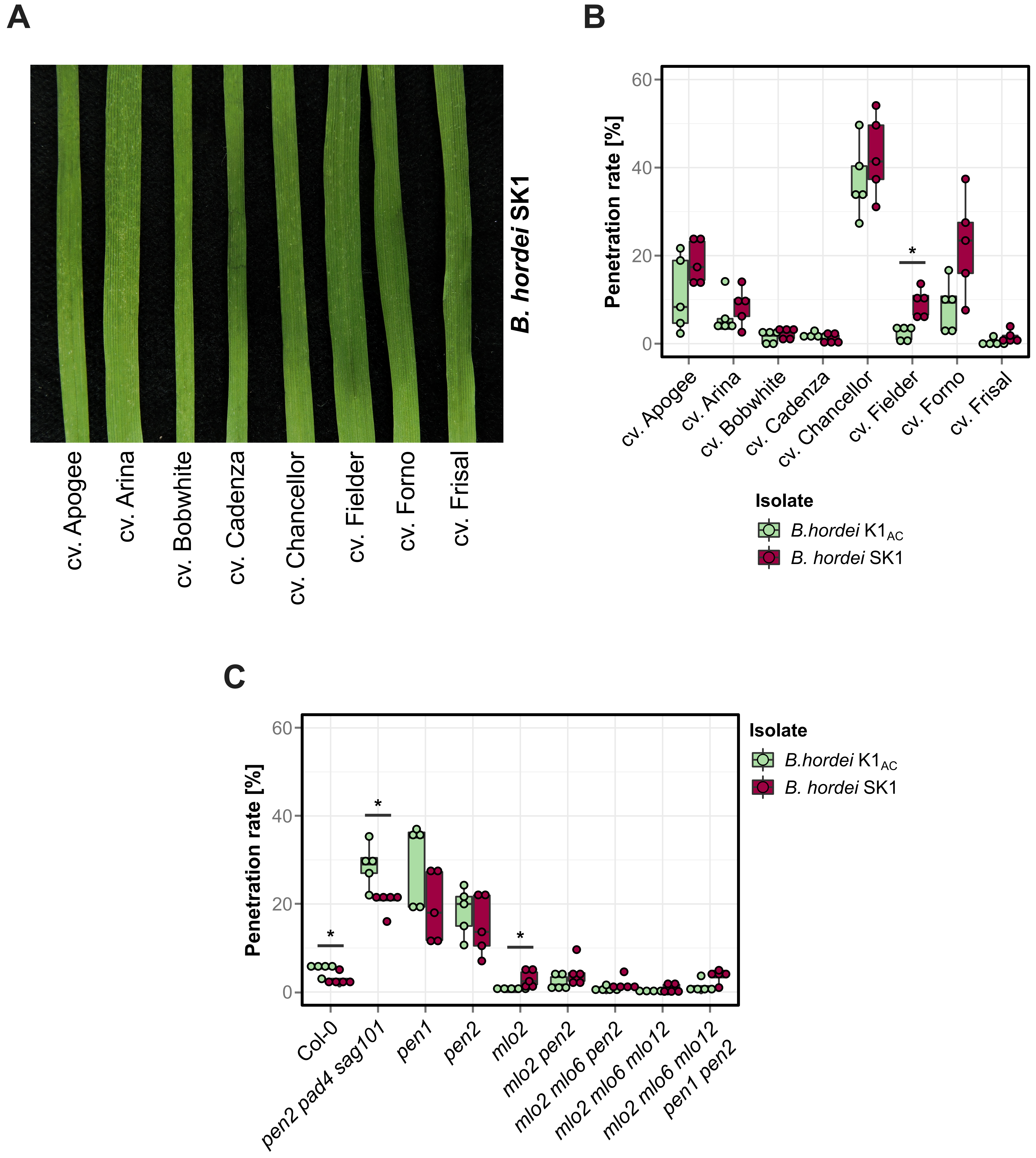

### Supplementary Figure 5

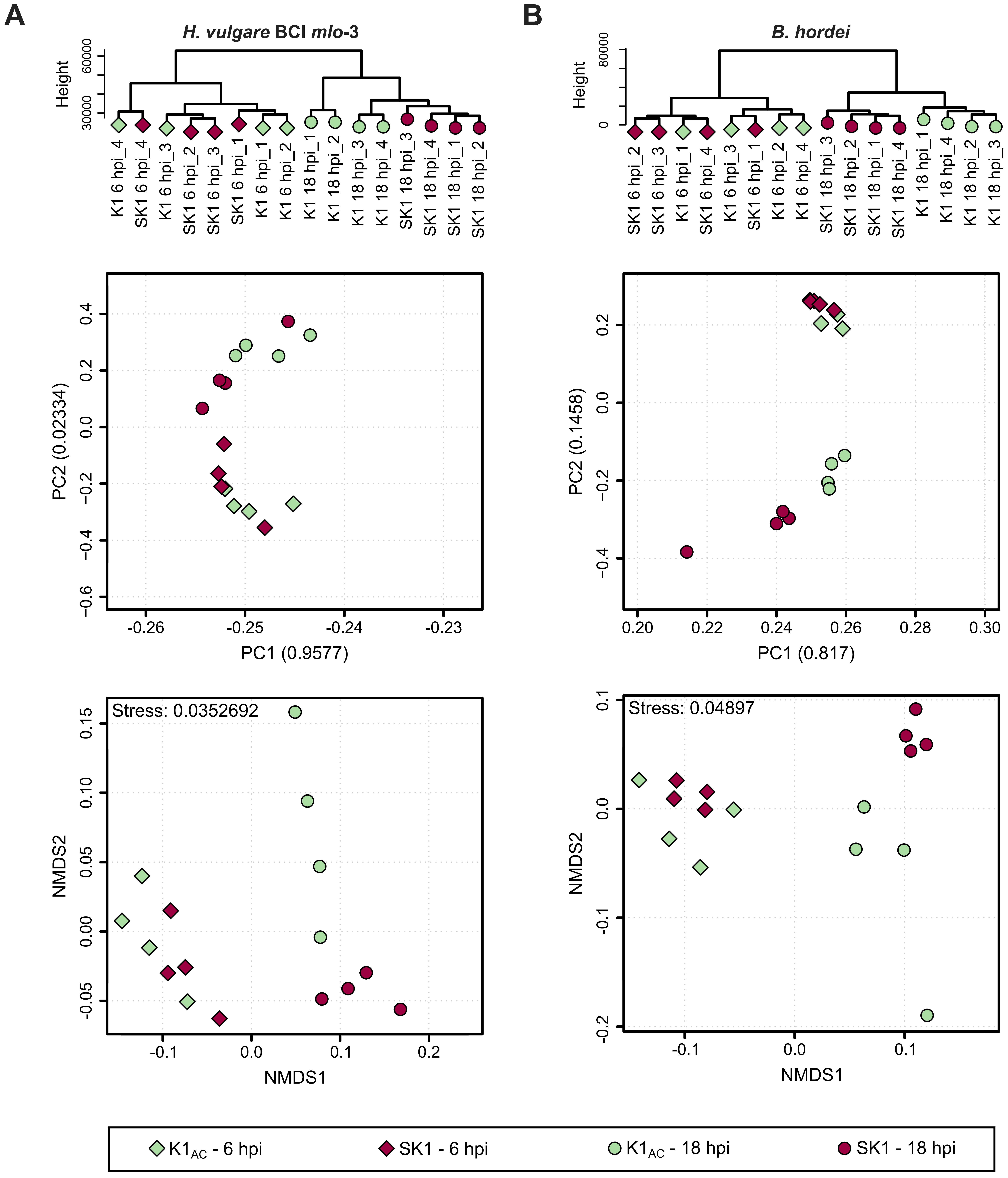

### Supplementary Figure 6

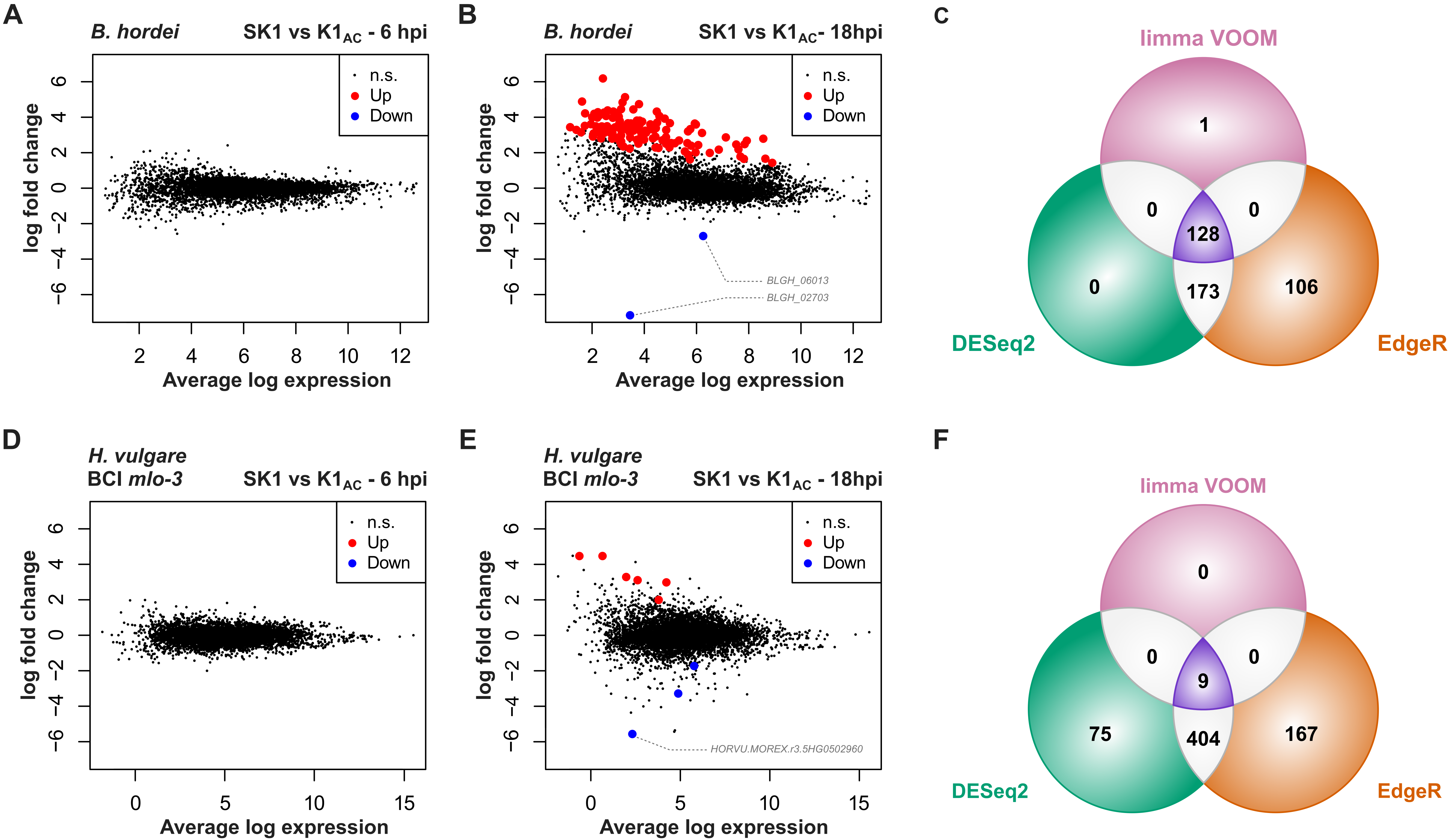

### Supplementary Figure 7

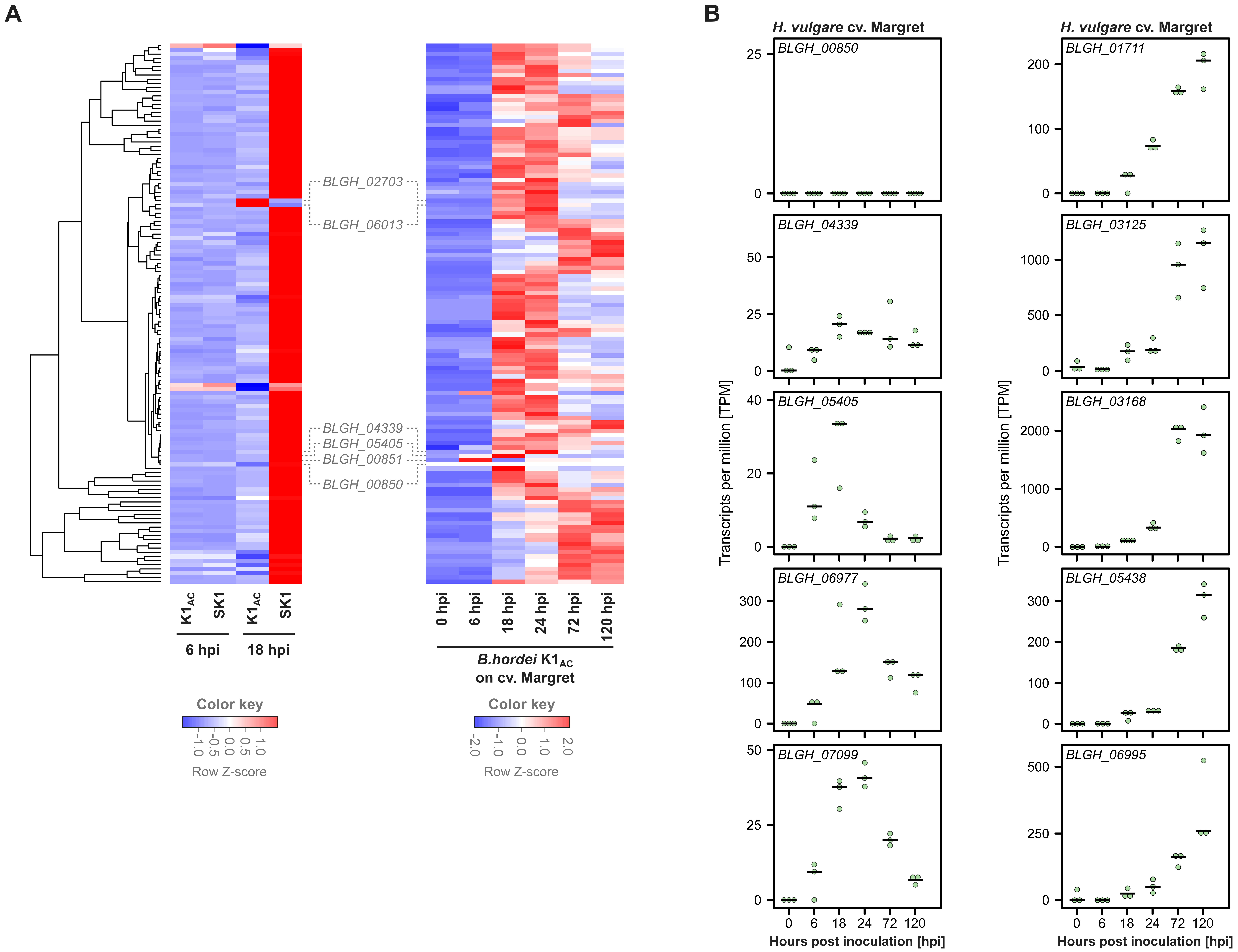

### Supplementary Figure 8

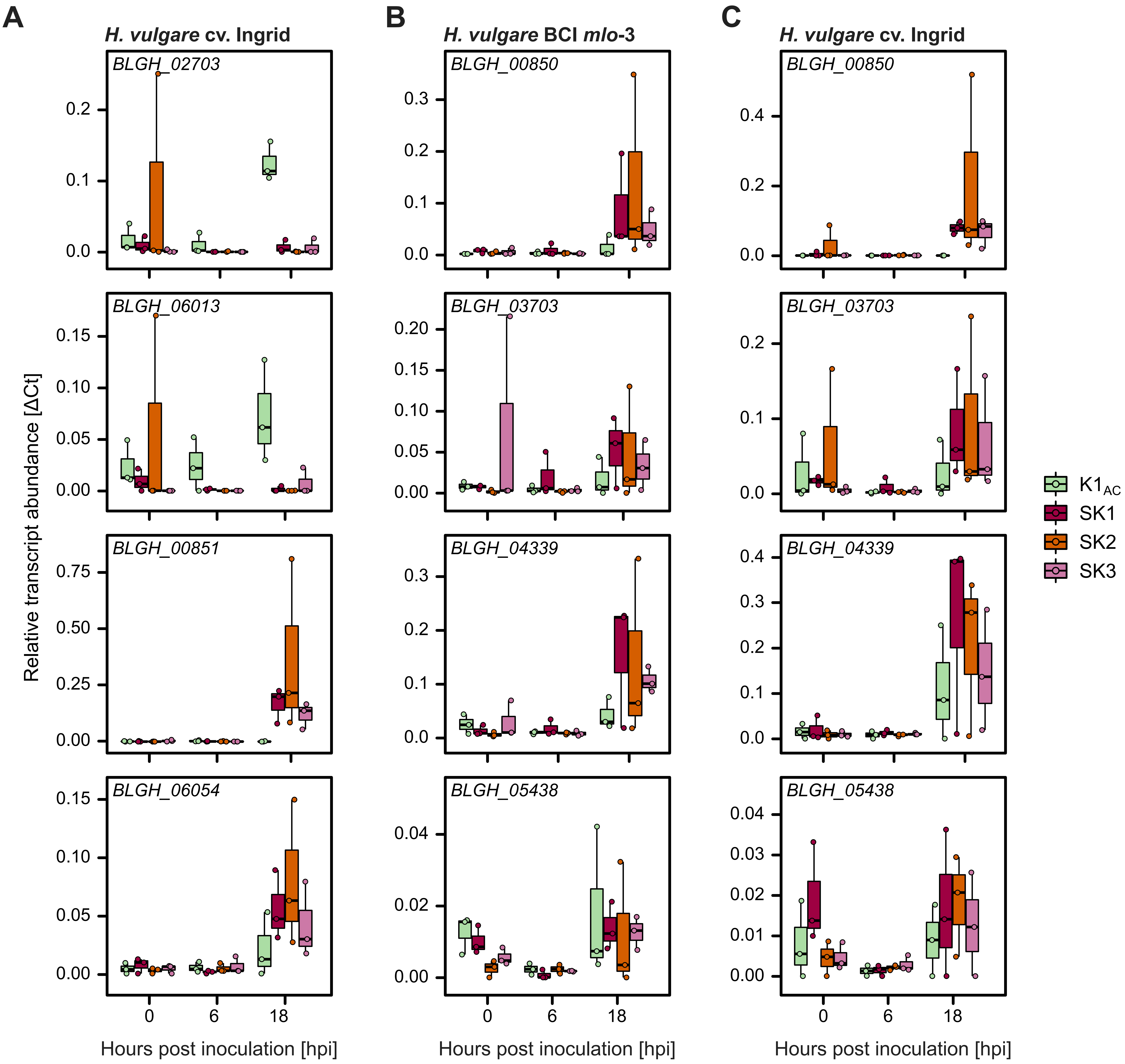

### Supplementary Figure 11

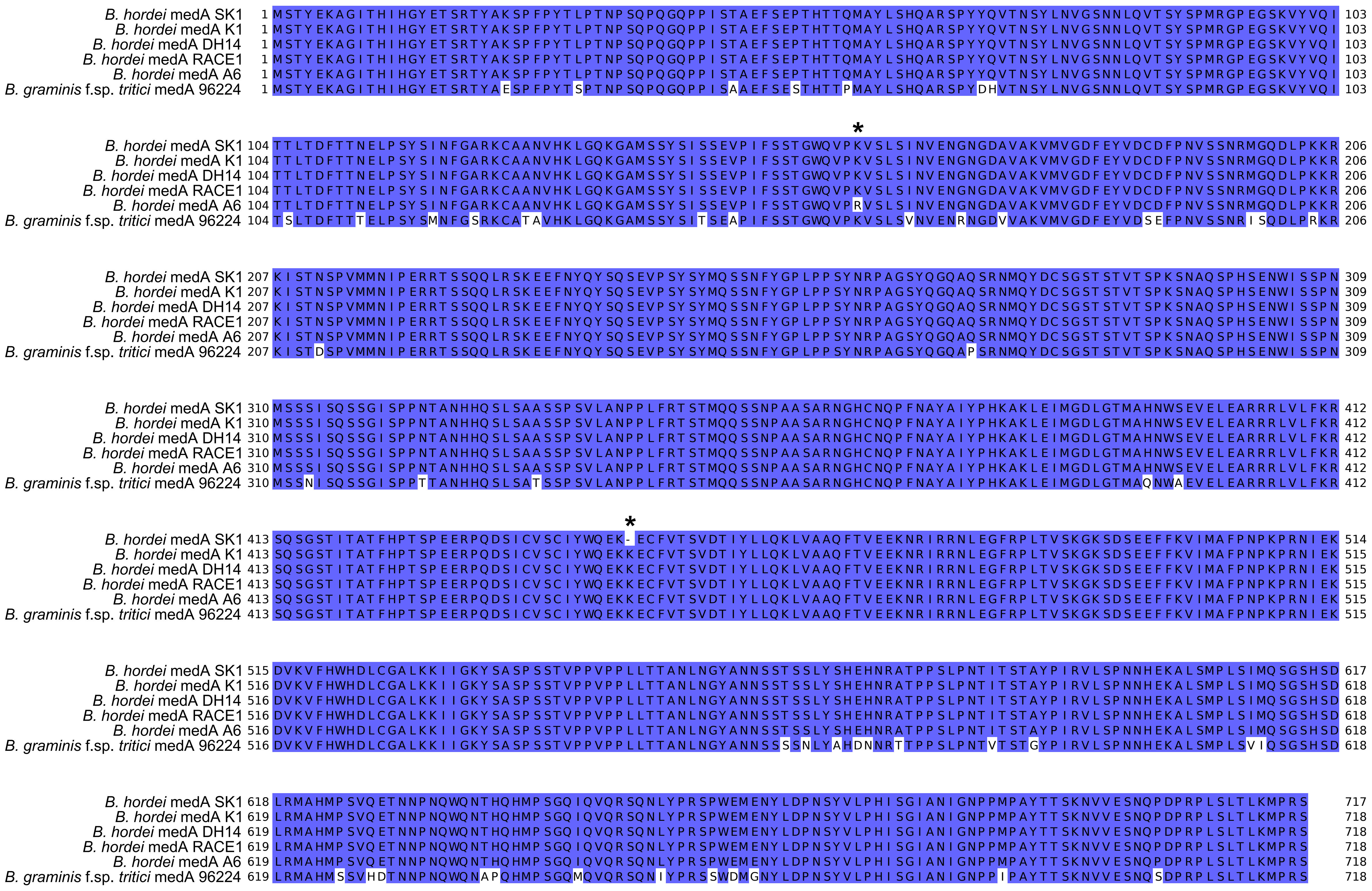

### Supplementary Figure 12

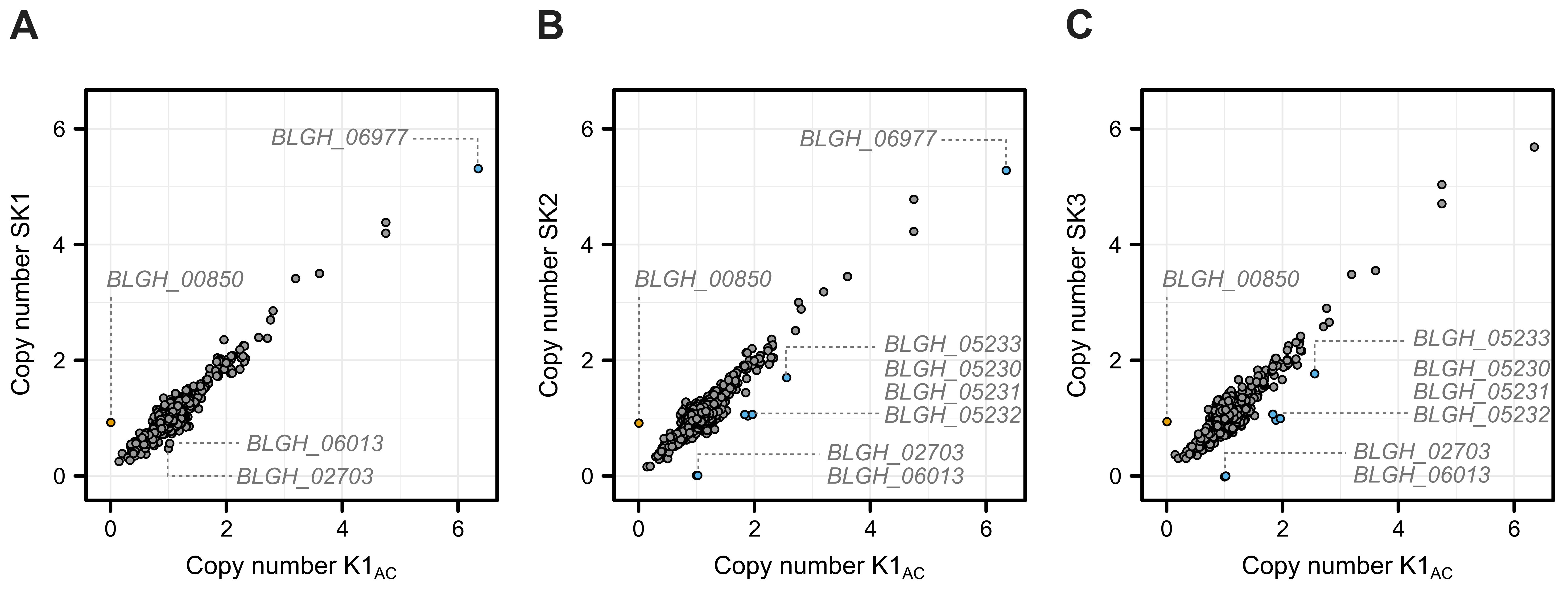

### Supplementary Figure 13

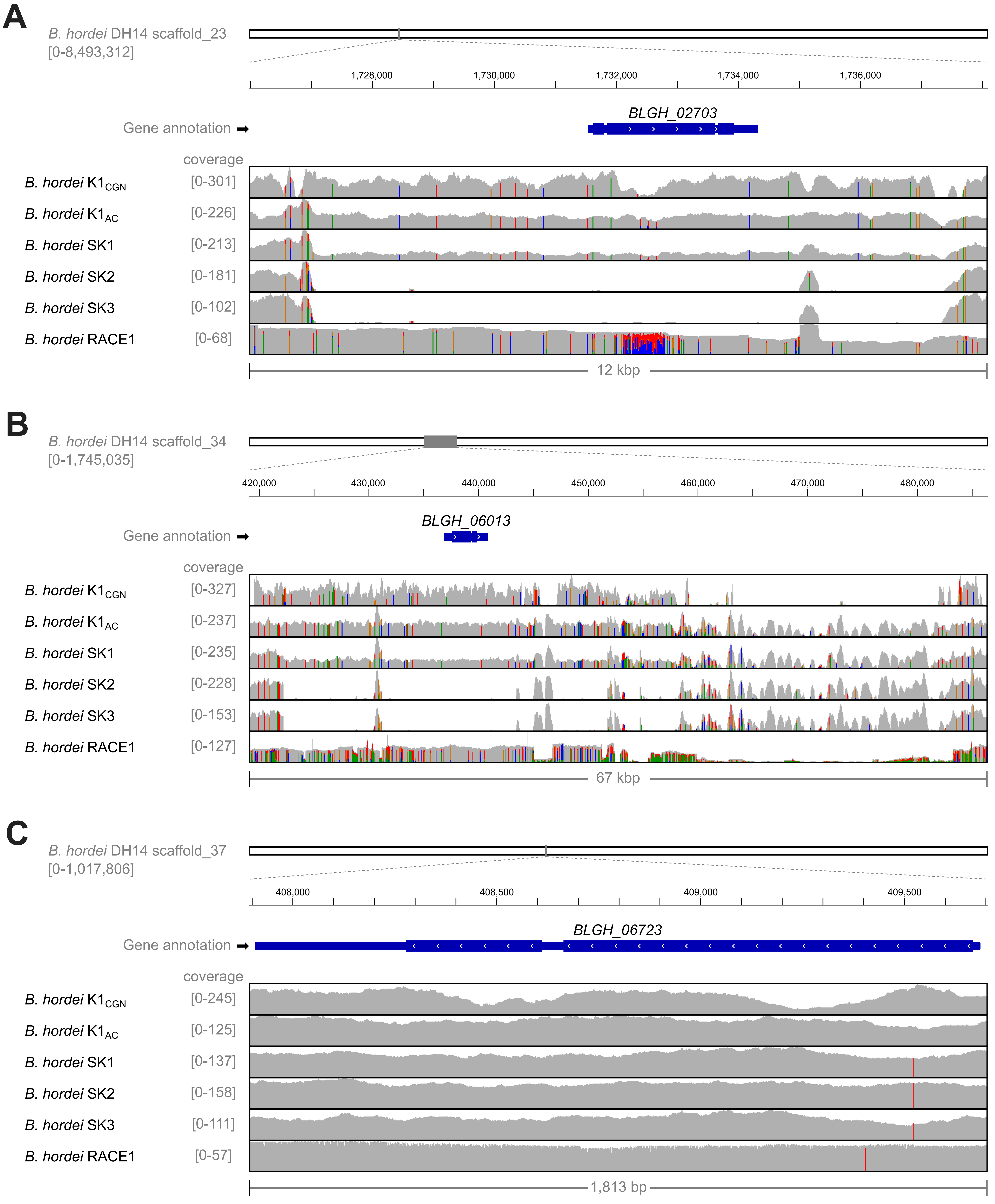

### Supplementary Figure 14

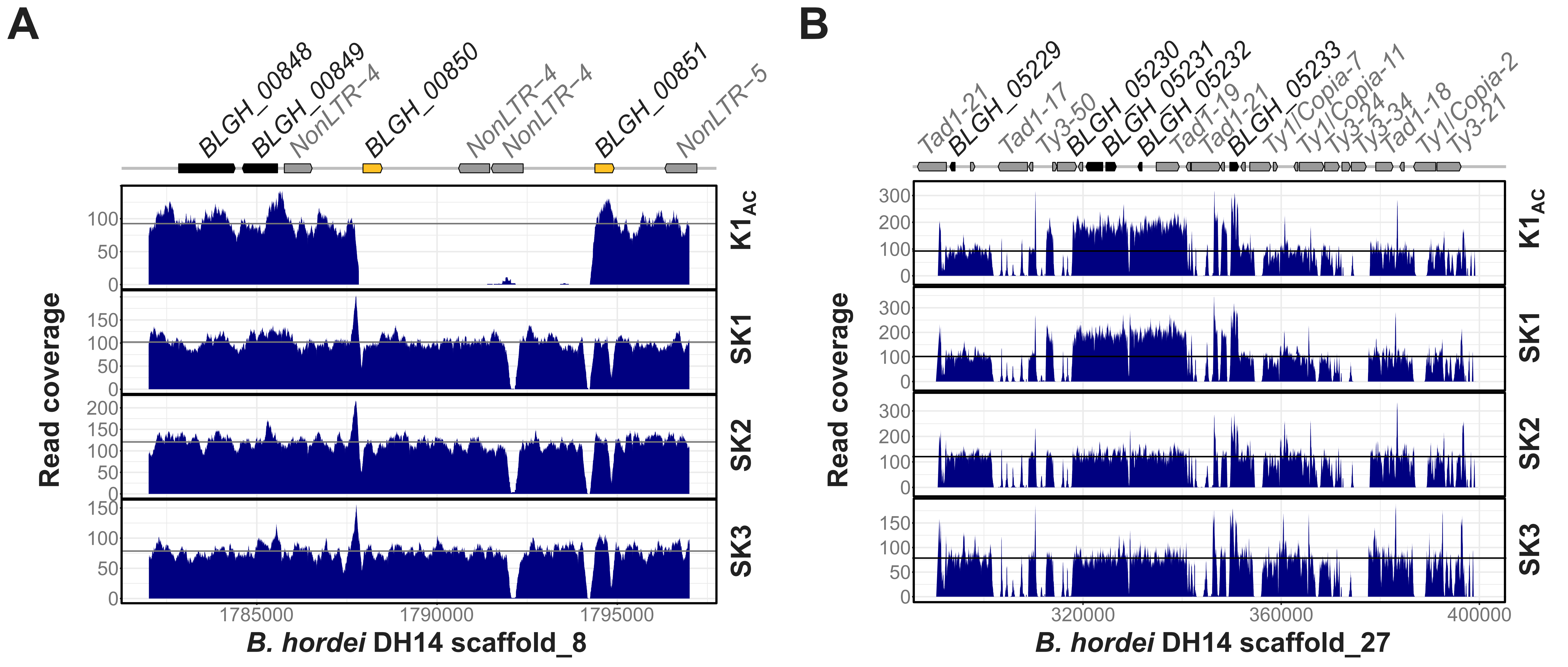

### Supplementary Figure 15

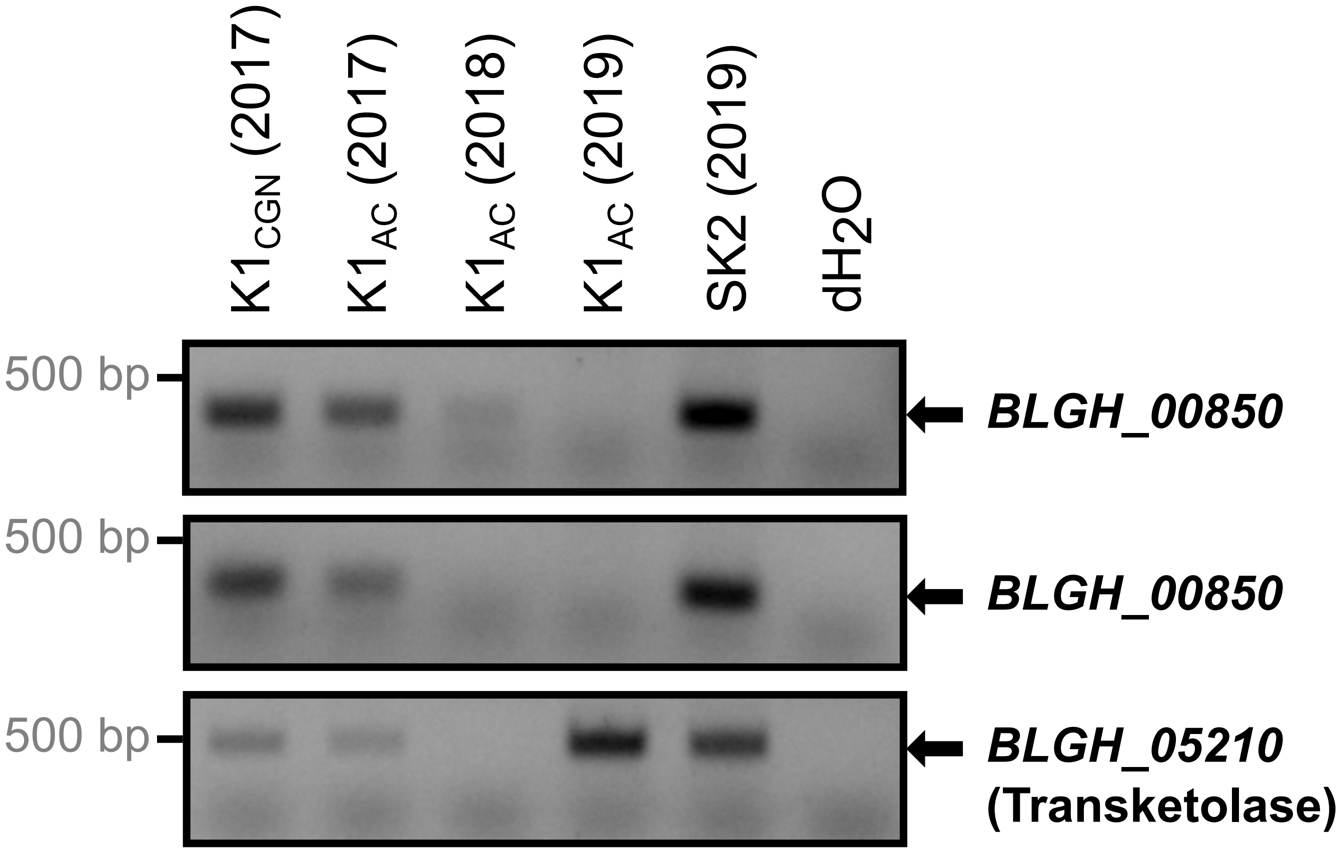

### Supplementary Figure 16

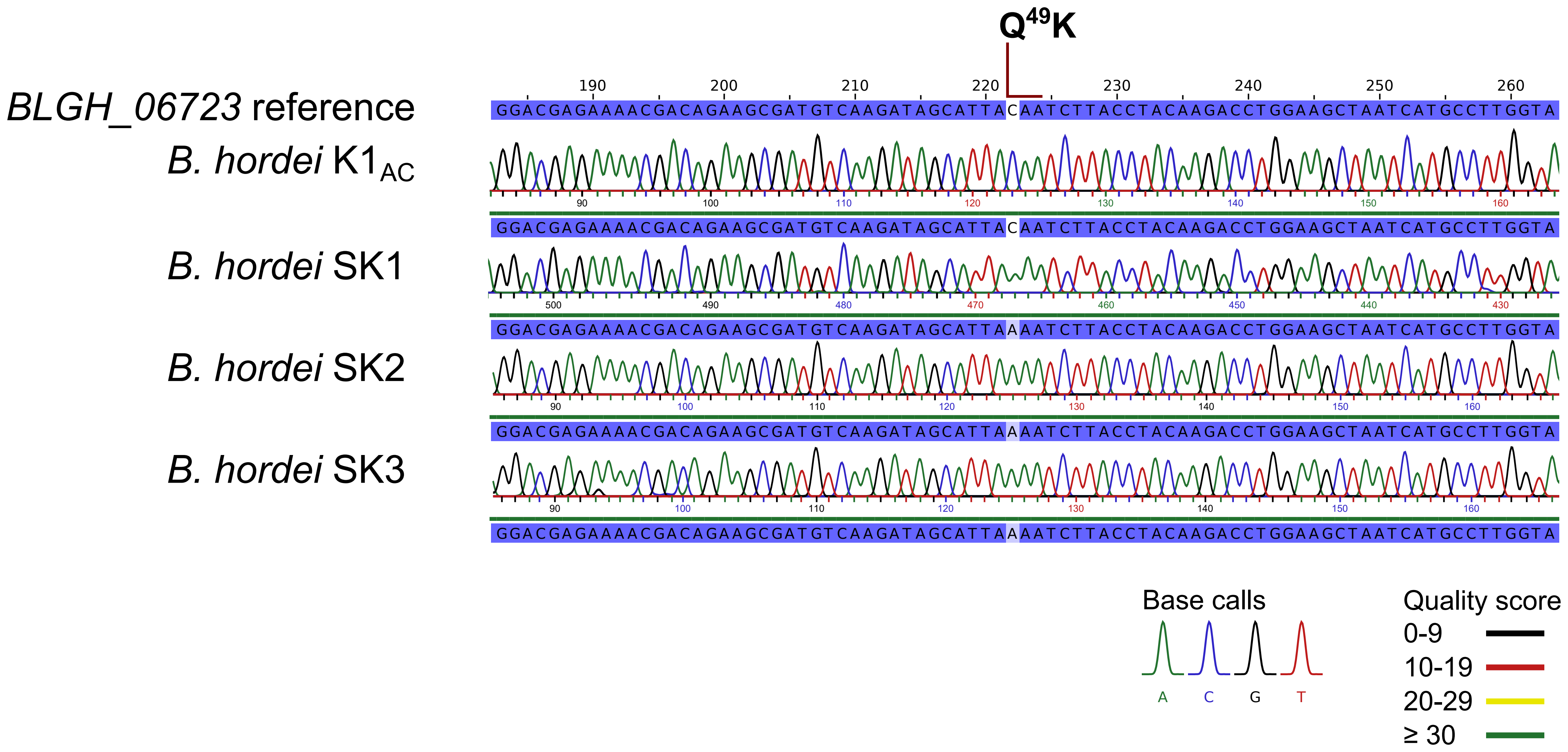

### Supplementary Figure 17

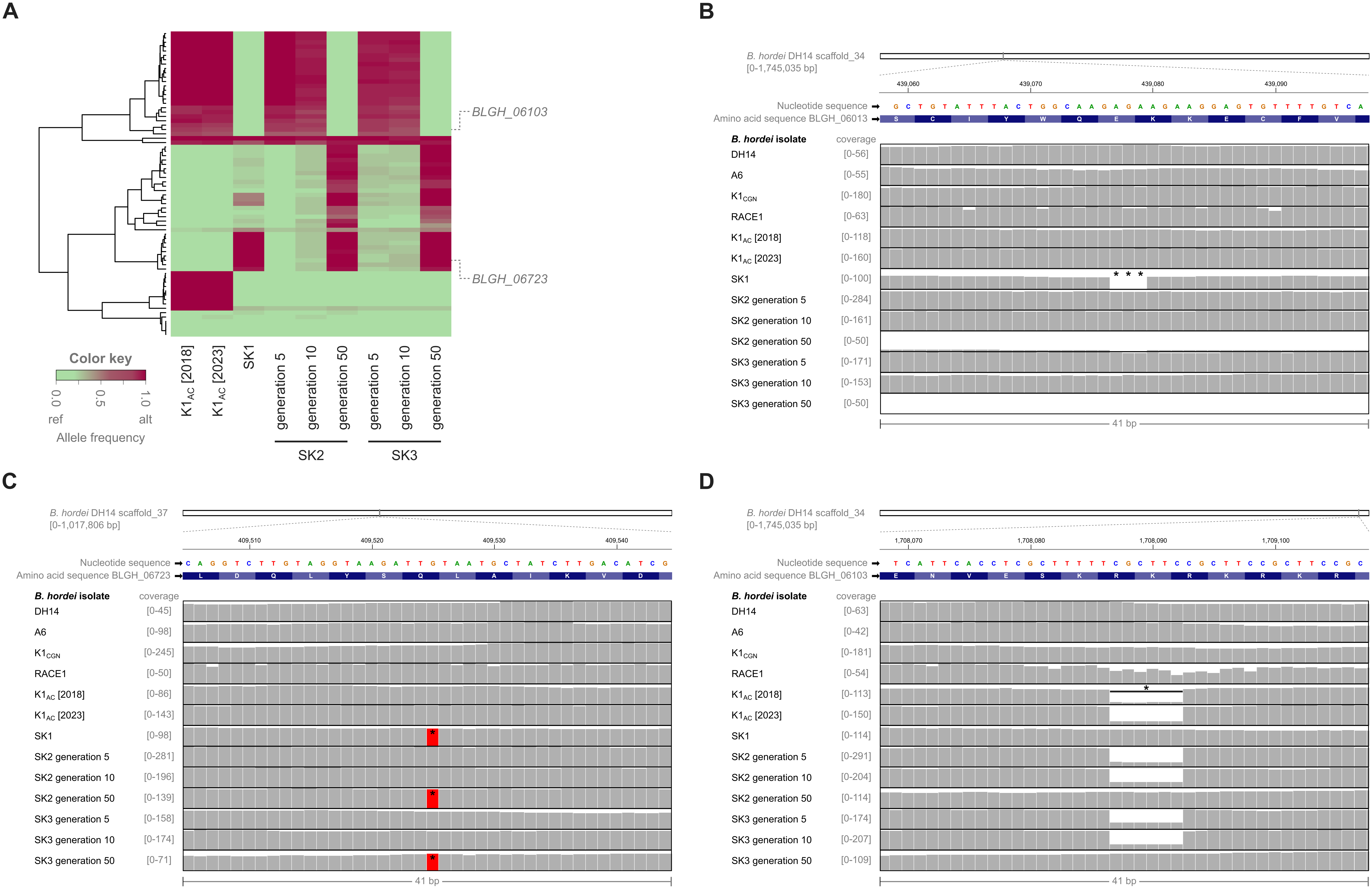

### Supplementary Figure 18

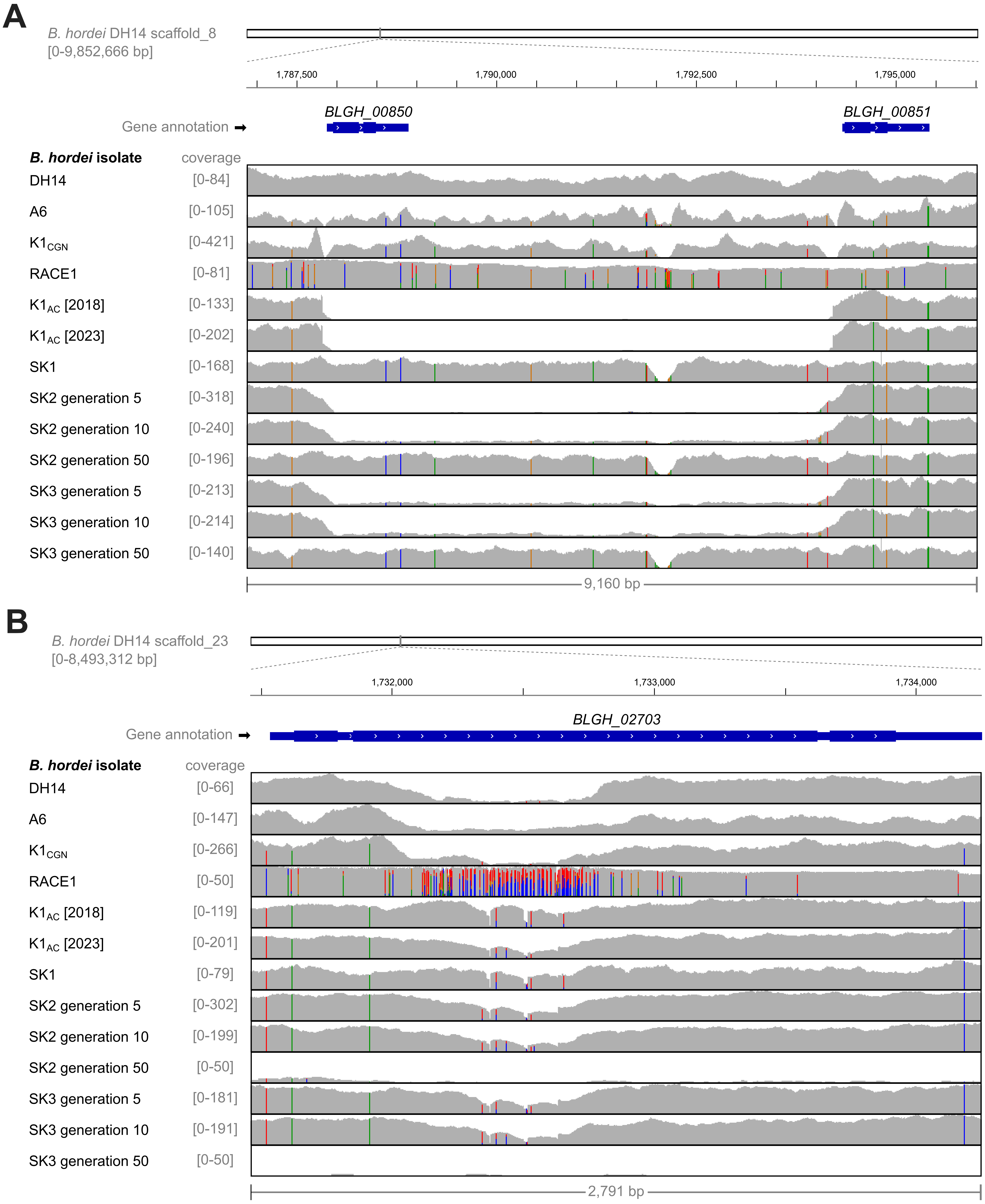

### Supplementary Figure 19

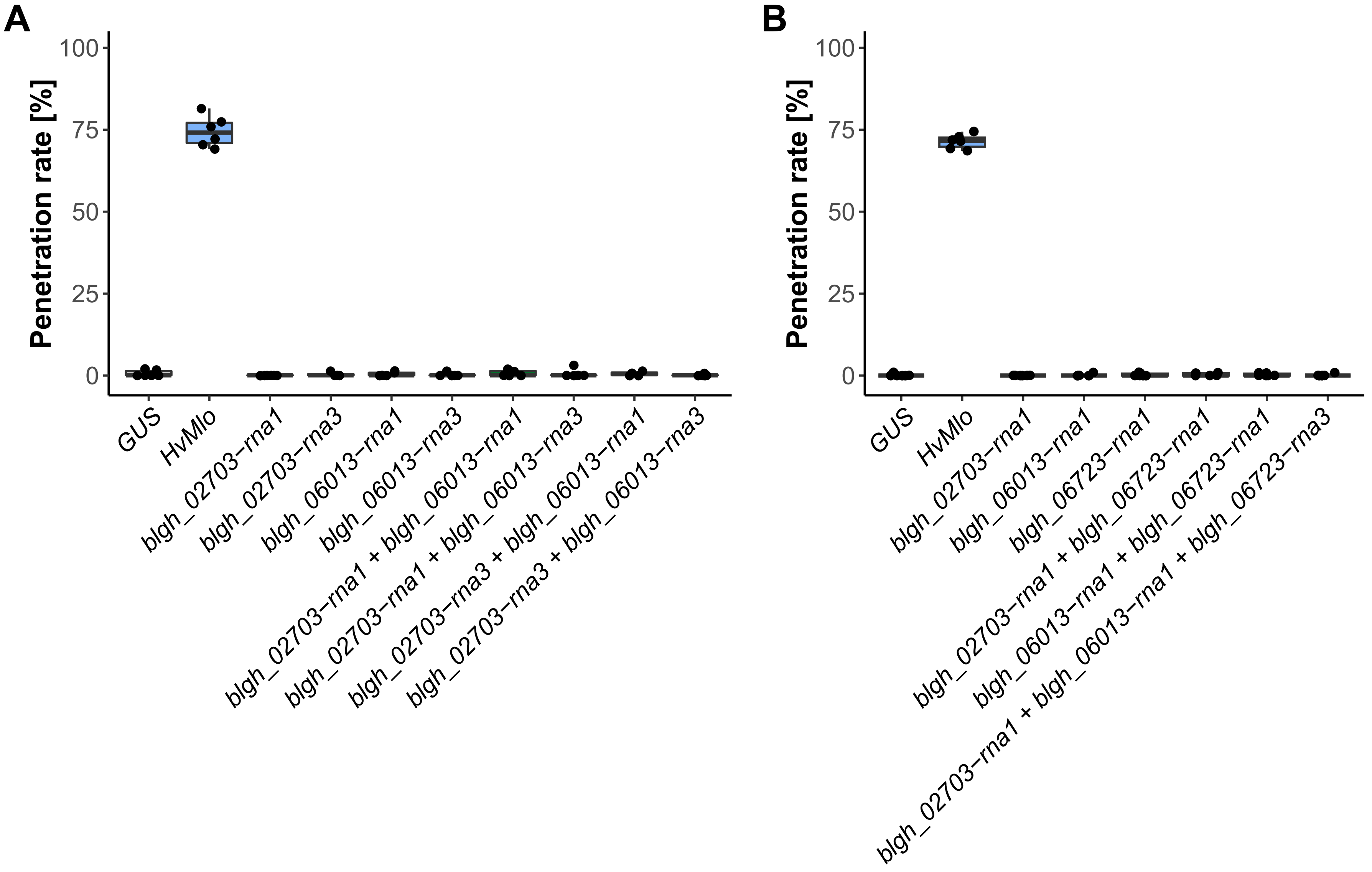

### Supplementary Figure 20

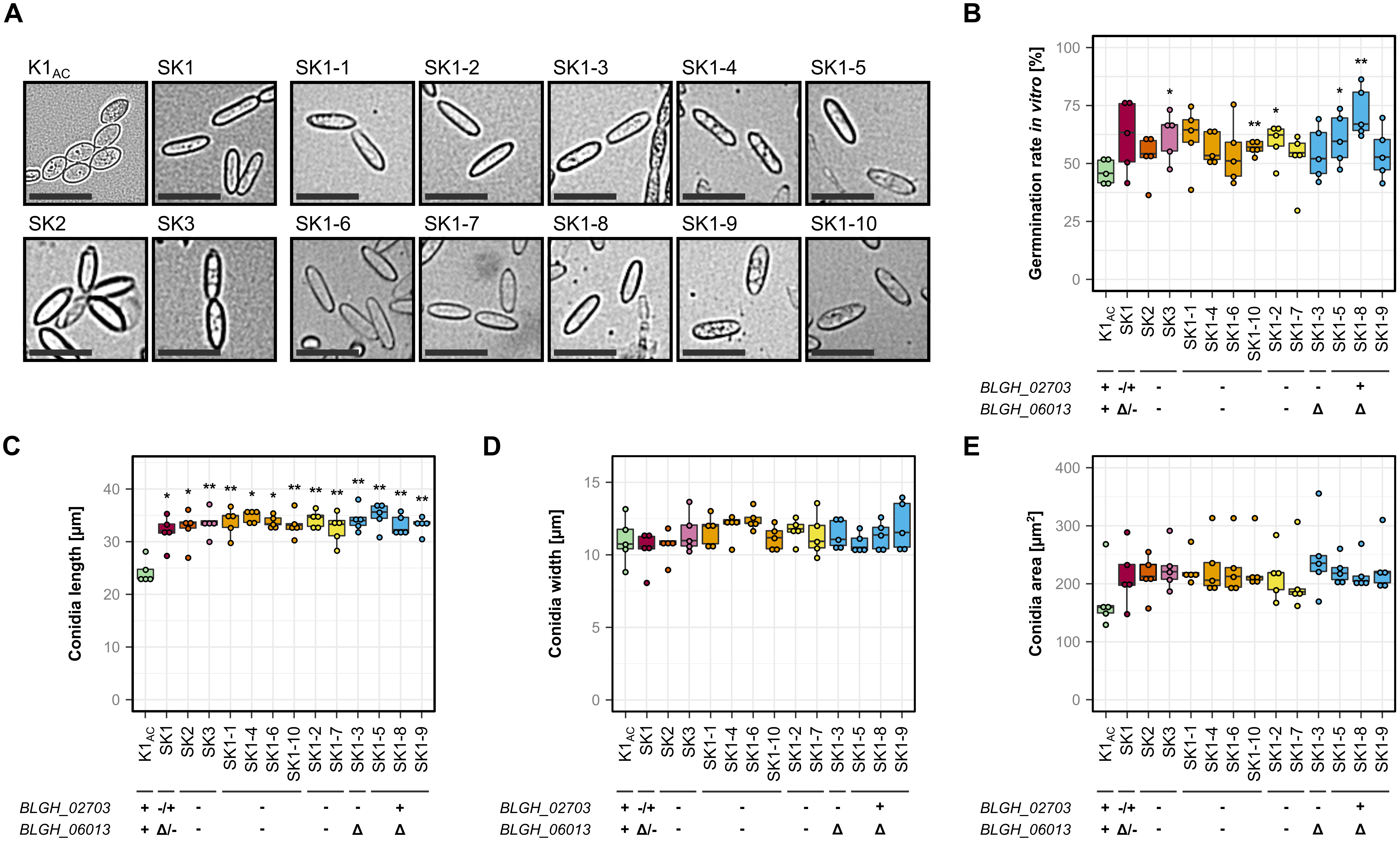

### Supplementary Figure 21

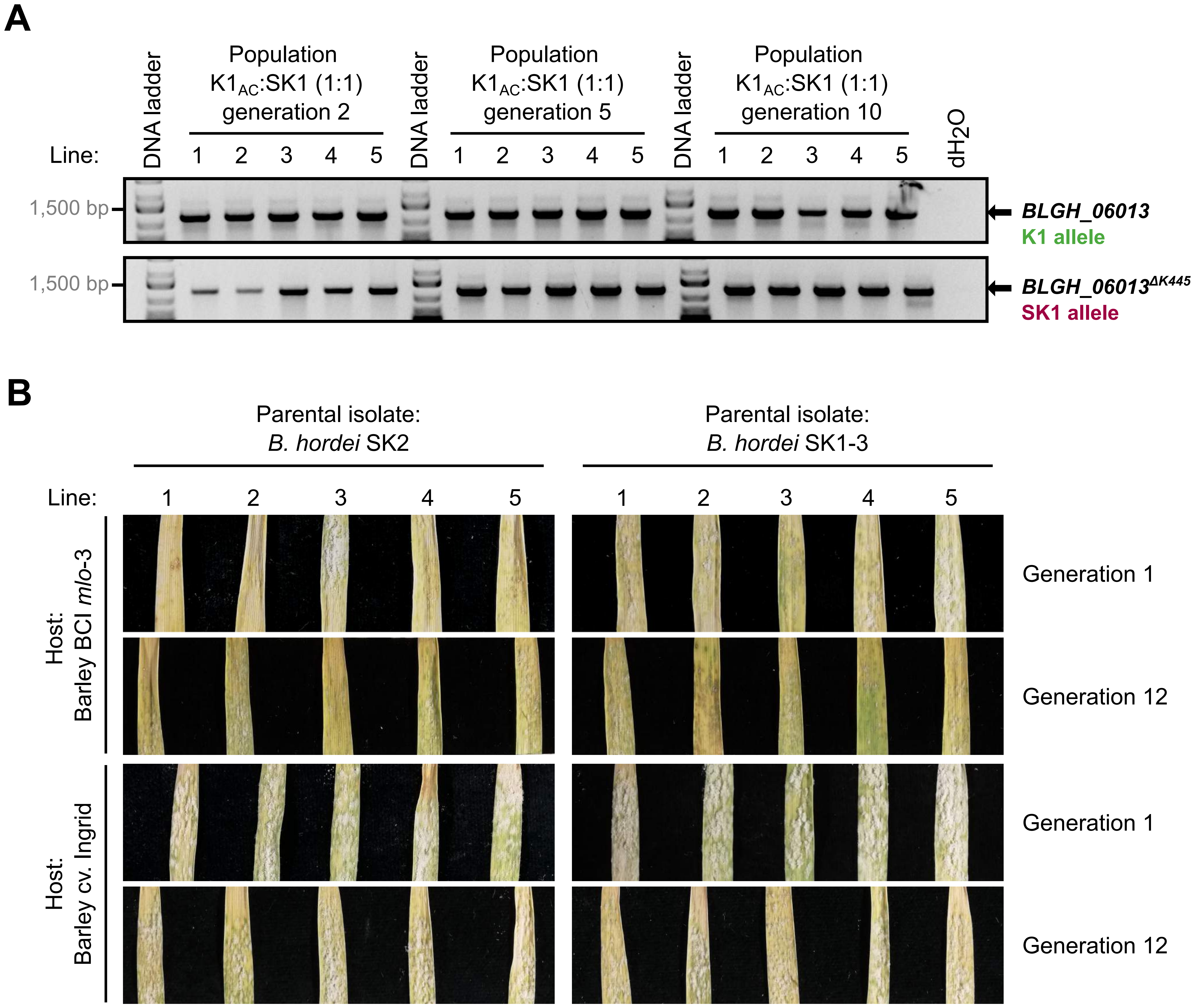

### Supplementary Figure 22

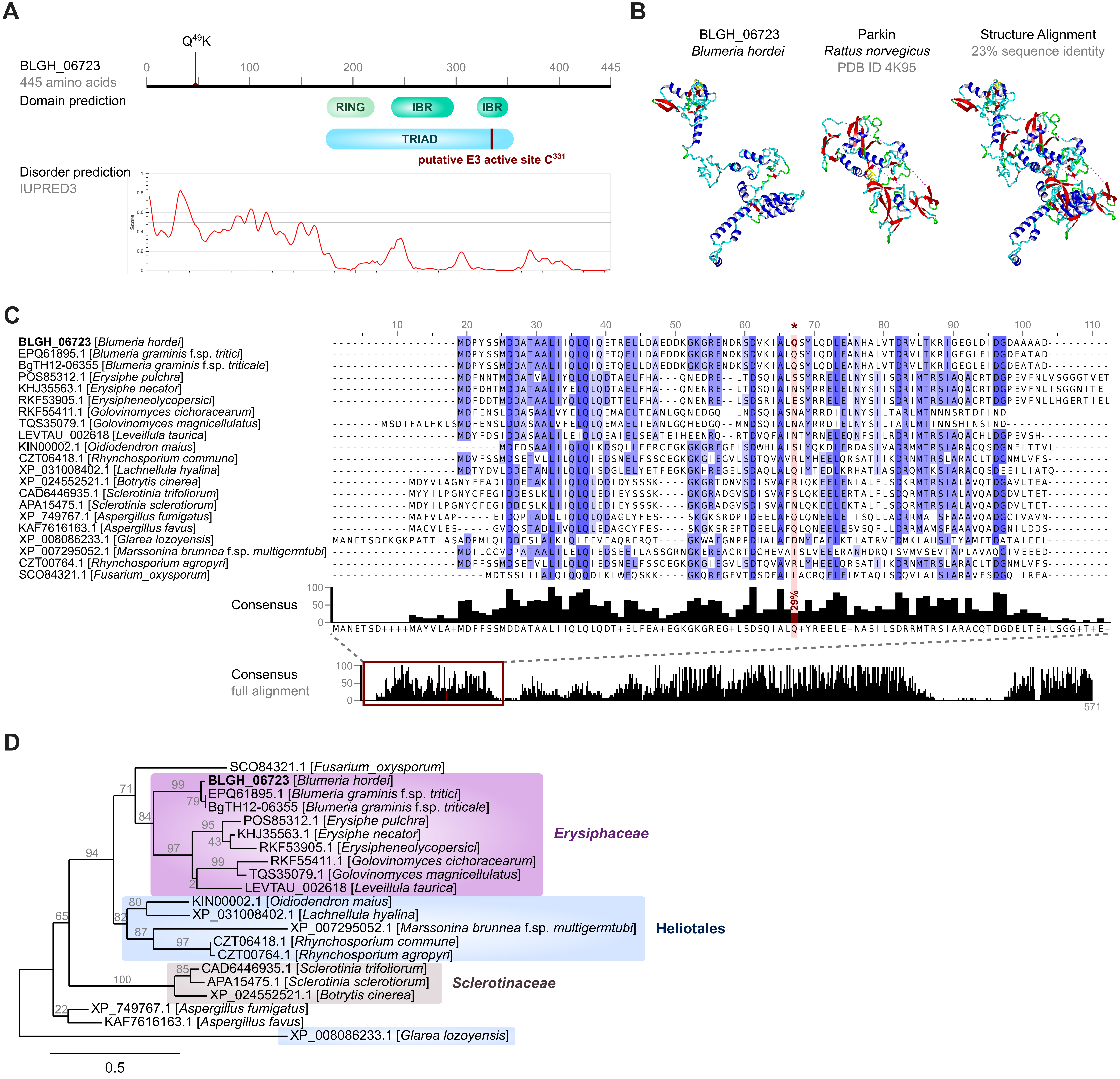
