## Supplementary Figure 9 for "A fungal plant pathogen overcomes *mlo*-mediated broad-spectrum disease resistance by rapid gene loss"

```
BLGH_06013     1 MSTYEKAGITHIHGYETSRTYAKSPFFPYTLPTNPSQPQGQPPISTAEFSEPTHTT  55
XP_755658.1   39 LSDYDSA--RSLHNGASYSTYGQTPYVTATSLVPS-PMADQASQISDCGPYLPNQ  90

BLGH_06013    56 QMAYLSHQARSPYYQVTNSYLNVGSNNLQVTSYSPMRGPEGSKVYVQITTLTDFT 110
XP_755658.1   91 EYASSYEESRSPMLGAESRQL---P---EVVSYNPQRGSEGTRVFVQIQCPHDLH 139

BLGH_06013   111 TNELPSYSINFGARKCAANVHKLGQKGAMSSYSISSEVPIFSSTGWQVPKVSLSI 165
XP_755658.1  140 SSPYATLYLVFGSKKCECLPHFLGLQGSSFQYALSVDTPPFLATGSPSLAVPLQI 194

BLGH_06013   166 NVENGNGDAVAKVMVGDFEYVDCDFPNVSSNRMGQDLPKKRKISTNSPVMMNIPE 220
XP_755658.1  195 IIDNHQGGSVNTLQVGVYTY------EQTSQDSPSDESRKRRIASYSENVAYRPI 243

BLGH_06013   221 RRTSSQQLRSKEEFNYQYSQSEVPSYSYMQSSNFYGPLPPSYNRPAGSYQGQAQS 275
XP_755658.1  244 RRAAGQQ---------GYEDGRATSYSPYLQ-----PL-PAMNGFLPPYQGNASP 283

BLGH_06013   276 RNMQYDCSGSTSTVTSPKSNAQSPHSENWISSPNMSSSISQSSGISPPNTANHHQ 330
XP_755658.1  284 RVAPAQYSAGSSASHS-SIRAPSPLTPAW--SPSI-VNVTSDSR-APAYAIAHSL 333

BLGH_06013   331 SLSAASSPSVLANPPLFRTSTMQQSSNPAASARNGHCNQPFNAYAIYPHKAKLEI 385
XP_755658.1  334 RQHKSSSPARPANPTLIRTSTLQQ-----ANG-LGH-SQSFNPYAMYPSKAVLKL 381

BLGH_06013   386 MGDLGTMAHNWSEVELEARRRLVLFKRSQSGSTITATFHPTSPEERPQDSICVSC 440
XP_755658.1  382 NGDLDSMAEGWTKEEREAKRRLVQFTRMQNGSTIHADFKPVSPDERAPNSICISC 436

BLGH_06013   441 IYWQEKKECFVTSVDTIYLLQKLVAAQFTVEEKNRIRRNLEGFRPLTVSKGKSDS 495
XP_755658.1  437 IYWEGKDECFVTSVDTIYLLESLVGVRFTVEEKNRIRRNLEGFRPLTVSKAKADS 491

BLGH_06013   496 EEFFKVIMAFPNPKPRNIEKDVKVFHWHDLCGALKKIIGKYSASPSSTVPPVP-P 549
XP_755658.1  492 EDFFKVIMGFPAPKPRNIEKDVKVFPWKILSHALKKIIGKYSASYSSTAGALPTP 546

BLGH_06013   550 LLTTANLNGYANNSSTSSLYSHEHNRATPPSLPNTITSTAYPIRVLSPNNHEKAL 604
XP_755658.1  547 ISSTFAGNGAASDSGT-----EPYAAASPQSVSDAAPSSSYCHSIATP-----AY 591

BLGH_06013   605 SMPLSIMQSGSH---S-DLRMAHMPSVQETNNPNQWQNTHQHMPSGQIQVQRSQN 655
XP_755658.1  592 SPPTG--SSGPHMTGTVELR-AVLPAVTQPYHSMASGYAYPTVCQQQGQLSLTAP 643

BLGH_06013   656 LYPRSPWEMENYL-DPNSYVLPHISGIANIG-NPPMPAYTT               694
XP_755658.1  644 V-SRPSWNPNSLAHQPSASAAPSYSYLAPLPYSLPDPSHGC               683
```

Predicted monopartite NLS (score >3.0) by NLSmapper
