## Supplementary Figure 10 for "A fungal plant pathogen overcomes *mlo*-mediated broad-spectrum disease resistance by rapid gene loss"

CLUSTAL FORMAT: MUSCLE (3.8) multiple sequence alignment

```
RAQ47439.1      ------------------------------------------------------------
XP_755658.      ------------------------------------------------------------
g19421.t1       ------------------------------------------------------------
XP_0072880      MHTTYTTTTCYYYHSAAAAAAAAAADLPVQHSPITLFSSFSFPRRNSQLARRPLVDLTVER
XP_0080841      ------------------------------------------------------------
APA12674.1      ----------------------------MLLTTCLYYTIFLDDNLFIQKVRPSGWLHIFW
XP_0245519      ------------------------------------------------------------
CZT12971.1      ------------------------------------------------------------
CZT42388.1      ------------------------------------------------------------
KIN00506.1      ------------------------------------------------------------
XP_0310090      ------------------------------------------------------------
BLGH_06013      ------------------------------------------------------------
EPQ64905.1      ------------------------------------------------------------
POS86580.1      ------------------------------------------------------------
RKF58170.1      ------------------------------------------------------------


RAQ47439.1      ------------------------------------------------------------
XP_755658.      ------------------------------------------------------------
g19421.t1       ------------------------------------------------------------
XP_0072880      EITPRSGSLPVGRESDIPHGDALFGDGQPLYTLGLLGTCHCHRDKSAWPISSTFHGRRRQ
XP_0080841      ------------------------------------------------------------
APA12674.1      QSAYPIEHPRNLKFFVLVPFSIVIDSLKYFILLVSCSRDFVQKIYAFALLYSSPTAKLAY
XP_0245519      ------------------------------------------------------------
CZT12971.1      ------------------------------------------------------------
CZT42388.1      ------------------------------------------------------------
KIN00506.1      ------------------------------------------------------------
XP_0310090      --------------MDQKRRICYSTTRFERLALARFEICLSPYCYRLFSKNQYLSLSQPS
BLGH_06013      ------------------------------------------------------------
EPQ64905.1      ------------------------------------------------------------
POS86580.1      --------------------------------------------------MGNSFEPKKA
RKF58170.1      ------------------------------------------------------------


RAQ47439.1      ----------------------------------------------------MSTLQKPP
XP_755658.      ----------------------------------------------------MSTFQKPP
g19421.t1       --------------------------------------------------MSMSTYGKLP
XP_0072880      DRIPPITHPTCVRSAPVIIDDSYASPETAAFHEEGYEGALVARVSSPRDLLPMSAYGKAQ
XP_0080841      ----------------------------------------------------MSAYGKPA
APA12674.1      STYTLFDSEHLSSGRPIIIDETYPSPETTALHEGGYEGALVPRADSPTDILSMSAYGKPQ
XP_0245519      ----------------------------------------------------MSAYGKPQ
CZT12971.1      ----------------------------------------------------MSAYGKSL
CZT42388.1      ----------------------------------------------------MSAYGKSL
KIN00506.1      ------QPEHLPSNRPIILDETYESPETAALHEEGYEGALVARSGSPSSLISMSAYGKQP
XP_0310090      SWHTLRTPEPLSSHRPIILDESYASPESAAFHEAGYEGALVARVDSPNDLLSMSAYGKPA
BLGH_06013      ----------------------------------------------------MSTYEKAG
EPQ64905.1      ----------------------------------------------------MSTYEKAG
POS86580.1      LSQTCPPNHLSTPNKTVILDDSDVSPETNTLHEVGNEEVYAIDKYLPKDRTSMSAYGKTP
RKF58170.1      ----------------------------------------------------MPAYERAP
                                                                    *.:   .

RAQ47439.1      QAALLD---------------------------YESARSVQDGASYSSFGQAPYVTSTSM
XP_755658.      SAALLGKFVPSFFLEHCCSFGKCRQTDRLTLSDYDSARSLHNGASYSTYGQTPYVTATSL
g19421.t1       TIHYHA---------------------------YDTGRFYAHPH--FQYVPQIYPSQSLD
XP_0072880      PPQLHG---------------------------YDAGRGYSDNT--FPYTAPSYGSQPPA
XP_0080841      PSELHS---------------------------YEPNRAYPEGS--FPYAT-TYGSQP-G
APA12674.1      PPQLHG---------------------------YETGRPYSENA--FPYSTPPYGTQSSI
XP_0245519      PPQLHG---------------------------YDTARSYPENA--FPYQTPSYGTQSSI
CZT12971.1      PPQLHG---------------------------FDGGRTYPENA--FPYAAPSYASQPPG
CZT42388.1      PPQLHG---------------------------YDGGRTYPENA--FPYAAPSYASQPPG
KIN00506.1      LPHLHG---------------------------FDAGRTFPDNS--FPYLSANYGSQPP-
XP_0310090      NTQSHG---------------------------YNTGRTYTEGP--FPYASPVYEAQSLS
BLGH_06013      ITHIHG---------------------------YETSRTYAKSP--FPYTLPTNPSQPQG
EPQ64905.1      ITHIHG---------------------------YETSRTYAESP--FPYTSPTNPSQPQG
POS86580.1      KTHNHEL--------------------------YEIGRSYPD-P--HSYTTTIYPSQSLG
RKF58170.1      STHSHV---------------------------YETGRAYADPP--YPYATNVCLSQSIG
                                                 ::  *          :      : .

RAQ47439.1      VPSPMADHASQISDCVP--YMGNQ-----EYASSYEDSRSPMLGGESRQLPEVISYSPQR
XP_755658.      VPSPMADQASQISDCGP--YLPNQ-----EYASSYEESRSPMLGAESRQLPEVVSYNPQR
g19421.t1       L-TIPPPGYSRLSHSTQLSYPAA------DARPTYDEIDGSYLNVGTS-TLELISFSPSK
XP_0072880      PAPTAAPLYQHAPQLTPVAYPPN------QVRTPYDEQTGPYLNVGVS-TPEITSYSPRR
XP_0080841      SSSATAPGYAHSQQLNPMAFPPS------QSRTPYDVQSGPYLNVGSTPIPEISKWEPKA
APA12674.1      PTSTPAPVYQNAGQMTSLGYASS------NTRTTYDEPAGPYLNVGSTPIPEVTSFSPQR
XP_0245519      PTSTPAPVYHNAGQMTPLGYGSG------STRTTYDEQSGPYLNVGSTPIPEVTSFSPQR
CZT12971.1      PATTTAPVYQTAPQLTPMTYTPN------QVRAPYDEQSGPYLNVGTT-IPEITSFAPNR
CZT42388.1      PATTTAPVYQTAPQLTPMTYTPN------QVRAPYDEQSGPYLNVGTT-IPEITSFAPNR
KIN00506.1      VPSSAAPSFSNIP-INPMGYAAD------QPRTSYDDQTGPYLNVGSTPTPEVASYSPSR
XP_0310090      VPASTAPGFQSTTQLTPMAYPANSVHAHTHAHVPYDDQSGPYLNVGSTPVPEVKSFSPHS
BLGH_06013      QPPISTAEFSEPTHTTQMAYLSH------QARSPYYQVTNSYLNVGSN-NLQVTSYSPMR
EPQ64905.1      QPPISAAEFSESTHTTPMAYLSH------QARSPYDHVTNSYLNVGSN-NLQVTSYSPMR
POS86580.1      LPSTIS-GYLGLSHSTSIPYSLN------ELRPAYHETPISYLDINSP-SPQLTSFSPSK
RKF58170.1      PPSIMS-AYLDMQPSTPITDFPS------EVRSIYHKNSGPFLDISGA-SPEVTSFSPSR
                  .  .                          *     . *.        :: .: *

RAQ47439.1      GSEGTRVFVQIQSPYDLHT-SSYATLYLVFGSKKCECVPHFLGFQ-GSLFQYALSVDTPP
XP_755658.      GSEGTRVFVQIQCPHDLHS-SPYATLYLVFGSKKCECLPHFLGLQ-GSSFQYALSVDTPP
g19421.t1       GPEYSKIHVDFT-FYEFIN-SNSLQFYLVLGSSKCAVVAQKLHPK-GILCSYRVSAEAPP
XP_0072880      GTRDTKFSVYISSLYELMSEDSTPHFFLMFGNVKCQAALQKLTQQ-GAVCSYSVTTEVPS
XP_0080841      GPASTPLHVYISSMYELMT-SNTPVFYLSFGGRKCQAALTKSSQQ-GAVCQYTVSAQVPD
APA12674.1      GSQGTTVSVYITTLYELLT-SNSPTFYLVFGQQKCQASVRKMNQS-GGVCSYVVTCEVPQ
XP_0245519      GSQGTQVSVYITTLYELLT-SNSPTFYLVFGQQKCQASVRKMNQS-GGVCSYIVTCEVPQ
CZT12971.1      GICGTKITVYITSLYELLT-DSAPVFSLMFGSRKCSASLQKVGQQ-GAVCSYTVVAEAPS
CZT42388.1      GICGTKITVYITSLYELLT-DSAPVFSLMFGSRKCSASLQKVGQQ-GAVCSYTVVAEAPS
KIN00506.1      GASGTKIFVYITTLYELIT-TNSPAFFLMFGQRKCSASITKASSQGGGVCQYQITAEAPE
XP_0310090      GSEDTKIYVSITSLYELMT-SSTPEFFLMFGHRKCQASLAKASQQ-GGVCQYTVTTEVPH
BLGH_06013      GPEGSKVYVQITTLTDFTT-NELPSYSINFGARKCAANVHKLGQK-GAMSSYSISSEVPI
EPQ64905.1      GPEGSKVYVQITSLTDFTT-TELPSYSMNFGSRKCATAVHKLGQK-GAMSSYSITSEAPI
POS86580.1      GSEYSTIHIEFTTVYDITT-TNIPGLYLTFGSCKRSASAHKIPPK-TSSHSYYIAADVPS
RKF58170.1      GPEHTKIFIEFTSLYDVRT-KTDTSISLSFGAIKRPIVARQYSQK-NNIFSYQVSIEVPL
                *   : . : :    :. .        : :*  *           .     .* :  :.*

RAQ47439.1      FISTGSPSFAV-PLQVAMNAQNECPSTTLQVGVYTYEN------------------ASHP
XP_755658.      FLATGSPSLAV-PLQIIIDNHQGGSVNTLQVGVYTYEQ------------------TSQD
g19421.t1       FSKTFSTSRKI-QVSLVIESSYDVVVNKMVVGEFEYTDQQD-----------HKNSVSKN
XP_0072880      YSATGWTSSAAVPISMWMESGDGDLIGKLEVGDFTFVD--------------G--LSATS
XP_0080841      FAVTGCTTSQV-PVNMLMESGDGDVIASVEVGNFGYVS--------------D--VQNDY
APA12674.1      FASTSWSSSRV-PVSMFME-GDGDVISKVDVGDFTFEN--------------------GP
XP_0245519      FASTNWSTSRV-PVSMFME-GDGDVISKVDVGDFTFES--------------------DV
CZT12971.1      FNVTGWHSPMV-PLCMLMESGDGDAIEKVDAGQFTFID--------------G--QNENS
CZT42388.1      FNVTGWHSPMV-PLCMLMESGDGDAIEKVDAGQFTFID--------------G--QNENS
KIN00506.1      FASTGWSSVQV-PVFMFMESGDGDMIAKVDVGPFTYVD--------------GELPDDGA
XP_0310090      FSATNWGSAQV-PIDMFMESGDGDVMGKVAVGKFTYLS--------------GSTQGGSE
BLGH_06013      FSSTGWQVPKV-SLSINVENGNGDAVAKVMVGDFEYVDCD------------FPNVSSNR
EPQ64905.1      FSSTGWQVPKV-SLSVNVENRNGDVVAKVMVGDFEYVDSE------------FPNVSSNR
POS86580.1      FSTTGWPSSVV-PVTMTLETKDGEIASKMFIGEFEYIDCHEKRRDRSNRNSTSNSGNNHP
RKF58170.1      FQSTSWHSFKV-PISLIIEAGDSNLISKMFVGQFEYTDQS------------NSLVTDRN
                :  *         : : ::        .:  * : : .

RAQ47439.1      SPSD-------ESRKRRIMSYSEEPMSG--KRTTGPPIQGKEQPPAYS----------YS
XP_755658.      SPSD-------ESRKRRIASYSENVAYRPIRRAAGQQ----------GYEDGRATS--YS
g19421.t1       LGQA-------ICKKRKLSVDLQELVNSPPKKSVCRDRSEKNES-ICHYAQVDSASS-YS
XP_0072880      GSQE-------NSRKRKLSIDSTGLMRSSSKRVSNHQLRPKEEYGTYGYAQSETPPS-YS
XP_0080841      GAQD-------GSRKRKMSDASPELIKSPAKRNSAQMLRPKEEYTAYGYPTGQNNSS-YS
APA12674.1      QSLE-------GTRKRKISTESAELMRSPAKRNS-QQLKPKEEF-PYSFSSSDGSPSSYS
XP_0245519      HSLE-------GTRKRKISTESAELMRSPPKRNS-QQLKPKEEF-PYAFSSSDGSPSSYS
CZT12971.1      TSQE-------TARKRKMSADSADM-RSPMKRSSSQHLRSKDEYSTYGYSQSEGAST-FS
CZT42388.1      TSQE-------TARKRKMSADSADM-RSPMKRSSSQHLRSKDEYSTYGYAQSEGAST-FS
KIN00506.1      TTQD-------ASRKRKLSTESAELMRSPVKRIPAQQLRAKEDYNSFTYATADGGP--YS
XP_0310090      TPQD-------LSRKRKISSESAEL-QSPIKRASIQVLRPKEEFSTYGYSAADGGS--YA
BLGH_06013      MGQD-------LPKKRKISTNSPVMMNIPERRTSSQQLRSKEEF-NYQYSQSEVPS--YS
EPQ64905.1      ISQD-------LPRKRKISTDSPVMMNIPERRTSSQQLRSKEEF-NYQYSQSEVPS--YS
POS86580.1      SSGT-------VSKKRKASGDILELMNSPRKKLPCHQPRPKEEYANYQYASTDSQSL-YS
RKF58170.1      TGQDFQSSYIEVPKKRKIAEKLPEFINSPRKKTSHQQSQEKNEYNHCQYTHLDPAPS-YS
                           .**.             ..                           ::

RAQ47439.1      PYLQS---------LPAMNGFVAP--YHPTSS----------PRVAPAQYSTV-NATPQS
XP_755658.      PYLQP---------LPAMNGFLPP--YQGNAS----------PRVAPAQYSAG-SSASHS
g19421.t1       --SPNSNCYGPVT-SLCGARQTGY--FPSQNS----------LRPAHLVYSGP-ESISSP
XP_0072880      PYLQSSSNYNHL--VPPYHRSTGS--YQGPPPQQQHQQQQQQPRHLAYGYQSS-TTASPP
XP_0080841      PYMQASQGYSNA--VPQYNRPLAG--YPSQPS-----------RNIGYGYSNA-SNASPP
APA12674.1      SYLQASSSYSNL--LPQYNRSSGG--YPAQAA----------PRNIGYGYGNS-STTSPP
XP_0245519      PYLQASSAYSNM--MPQYNR-SGA--YSTQSA----------PRNMGYGYGSS-STASPP
CZT12971.1      PYLQPSNNYNNL--VPQYNRSVSG--YQGQQA-------------RNFGYYAT-STVSPP
CZT42388.1      PYLQPSNNYNNL--VPQYNRSVSG--YQGQQA-------------RNFGYYAT-STVSPP
KIN00506.1      PYMQQ-NAYANQ--LSQYGRAAPG--YQS-------------SRHLAFSYPSS-TASSSP
XP_0310090      PYLQPSASYGNL--LPQYNRSVGG--YQTQPS----------TRHLAYSYSNS-ATASPP
BLGH_06013      -YMQSSNFYGPL--PPSYNRPAGS--YQGQAQ----------SRNMQYDCSGSTSTVTSP
EPQ64905.1      -YMQSSNFYGPL--PPSYNRPAGS--YQGQAP----------SRNMQYDCSGSTSTVTSP
POS86580.1      -YLPNSNFYMPP--LTPYSGLQSSNFSGSQAS----------TKPIQYESLGS-RTAASP
RKF58170.1      -YLTASNFYGSTLGPPSSSDQEGN--FQVQPS----------LRSHLYGHTSS-ASTITA
                                                                           .

RAQ47439.1      NIRAPSPLAPSWSPSYVTVHGD-SRAPGFA-YGHGLRQ-----QKPTSPARLSNPTLIRT
XP_755658.      SIRAPSPLTPAWSPSIVNVTSD-SRAPAYA-IAHSLRQ-----HKSSSPARPANPTLIRT
g19421.t1       TLKSQSLPSGLWGSH-QNPSTNFSRSPGLV-SNSATHLTISSGSTASSPSTSQNLPLYRI
XP_0072880      TIKEQSPQASSWYPS----GPTAPRSQGAQ-SHLGPSR--PALSLL--PTSQTTPTLIRT
XP_0080841      TIKAQSPRVGSWSPAYGSAGQHLGRGSGLP-SSESMSR-----SSLPSPGLSANPPLIRT
APA12674.1      TLKAHSPHVSTWNSGYSTIGSSMARSPGL--PSNSMQR--SNMSSLPSPG---NPALIRT
XP_0245519      TLKAHSPHVSTWNSGYATIGSSMARSPGL--PSNSMQR--SNMTSLPSPG---NPALIRT
CZT12971.1      AIKAHSPQSANWSYQ----SASLARSSGIQ-TSLGLQR--PGLVSL--PASSANPPLIRT
CZT42388.1      AIKAHSPQSANWSYQ----SASLARSSGIQ-TSLGLQR--PGLVSL--PASSANPPLIRT
KIN00506.1      TVKAHSPQSSTWNPSYSNLGSGMARGPGE--PPGGSHR--GSVTSLPSPAIPANPPLIRT
XP_0310090      TMKAQSPQVGSWSPAYGAVSSNMVRSPGIP-SSGGVTR--PSLSALPSPAASANPRLIRT
BLGH_06013      KSNAQSPHSENWISS-PNMSSSISQSSGISPPNTANHH--QSLSAASSPSVLANPPLFRT
EPQ64905.1      KSNAQSPHSENWISS-PNMSSNISQSSGISPPTTANHH--QSLSATSSPSVLANPPLFRT
POS86580.1      TDKGTSPQPRKWPSN-SNLRQNVKQSPAMS-PNSTSHQ--GSVSSNSSPSVLANPPLFRT
RKF58170.1      TPKDQSSQCGNWNFQ-MNLGSNISQSPDTS-PKSASHH--LS-TPASLPSALANPPLFRT
                  .  *     *           ..                       *     .  * *

RAQ47439.1      STLQQ-------SSSLGHHTQSFNPYA--MYPSKAVLKLNGSLDS-MTESWTKEEREAKR
XP_755658.      STLQQ-------ANGLGHSQS-FNPYA--MYPSKAVLKLNGDLDS-MAEGWTKEEREAKR
g19421.t1       STYQQ-SSCPVGNPHLQQSKQQFNAYS--LYPNKAKVEIVSDLAT-MTQGWTDQEIKNRR
XP_0072880      STIQQ-TPSPATTPHGGAH---FSSYGGLMYPHKAKLEIHGNLDD-MTTGWSDDEWESRR
XP_0080841      STLQQ-TPSPATIPHGGIHGQQFAPYN--LYPSKAKLEINGDIES-MAKEWSEDEWDAKR
APA12674.1      STLQ--SSSPGATPHAGHT---YNPYP--MYANKAKLEISGDLGDMMVNNWTAEEWESKR
XP_0245519      STLQ--SSSPGATPHAGHT---YNPYP--MYANKAKLEISGDLGDMMVNNWTAEEWESKR
CZT12971.1      STLQQNSPSPATTPHGGHPAH-FNAYA--LYSNKAKLEVAGDLEE-MPVHWSSEEFDSRR
CZT42388.1      STLQQNSPSPATTPHGGHPAH-FNAYA--LYSNKAKLEVAGDLEE-MPVHWSNEEFDSRR
KIN00506.1      SILQQ-PPSPATTPHGAHAGQQFNAYA--LYPHKAKLELQGDLDA-MARNWTEEEWEQKR
XP_0310090      STLQQ-TPIPASTPHAAHPGQQFNAYA--LYPHKAKLEINGDLDA-MAQSWSEEEWDLKR
BLGH_06013      STMQQ-SSNPAASARNGHCNQPFNAYA--IYPHKAKLEIMGDLGT-MAHNWSEVELEARR
EPQ64905.1      STMQQ-SSNPAASARNGHCNQPFNAYA--IYPHKAKLEIMGDLGT-MAQNWAEVELEARR
POS86580.1      STFHQ-NSSPADNHQLRNSKQ-FNSYS--LYPHKAKLEIIGDLNS-MAQSWTDEEINFRR
RKF58170.1      STLQQ-NFPPVDTSCPEQAKQ-FNTYT--MYPNKAILEIIGELGS-MTKNWKNEEIKCRR
                *  :                   : .*   :*. ** ::: ..:   *    *   * . .*

RAQ47439.1      RLVQFTRMQSGSTIHADFNPVAPEDRA---PNSICISCIYWDGKDECFITSVDTIYLLES
XP_755658.      RLVQFTRMQNGSTIHADFKPVSPDERA---PNSICISCIYWEGKDECFVTSVDTIYLLES
g19421.t1       RLVYFSRVQSGSTITTDFRPLLPTDHP---EPGSCVSCIHWKEKKDCFVTSVDTIQLLQF
XP_0072880      RLVHFRQAQSGSTITVTFTPVTPEDRPQSTPTSGTISCIYWEEKNECFVTSVDTICLLEK
XP_0080841      RLVVFTRSQSGSTISASFRAVTPEERP---PQSICISCIYWEEKEQCFVTSVDTISLLEQ
APA12674.1      RLVQFKRSQSGCTITTTFNSISADERP---PQSVVISCIYWEEKNDCYVTSVDTITLLEA
XP_0245519      RLVQFKRSQSGCTITTTFNAISADERP---PQSVVISCIYWEEKNDCYVTSVDTITLLES
CZT12971.1      RLVHFRRSQSGSTITASFNAVSQEERP---QNSVCISCIYWEEKQECFVTSVDTIYLLEH
CZT42388.1      RLVHFRRSQSGSTITASFNAVSQEERP---QNSVCISCIYWEEKQECFVTSVDTIYLLEH
KIN00506.1      RLVHFKRSQSGSTITTQFQPVTLEDRP---QGSICISCIYWEERNECFVTSVDTIFLLEQ
XP_0310090      RLVHFRRSQAGSTITTTFQPVSVDERP---QDSVCISCIYWEEKQDCFVTSVDTIYLLER
BLGH_06013      RLVLFKRSQSGSTITATFHPTSPEERP---QDSICVSCIYWQEKKECFVTSVDTIYLLQK
EPQ64905.1      RLVLFKRSQSGSTITATFHPTSPEERP---QDSICVSCIYWQEKKECFVTSVDTIYLLQK
POS86580.1      RLVLFKRSQSGSTITTKFKSVRPDDRP---KDSICISCIYWEEKRDFFITSVDTIYLLQK
RKF58170.1      RIVVFKRSQVGSTITASFHPLGPDERP---TDNICISCIYWEEKNDYFVTSVDTIFLLQK
                *:* * . * *.** . * .    :..     .  :***:*. . : ::****** **:

RAQ47439.1      LVGVRFTVEEKNRIRRNLEGFRPLTVSKAKADSEDFFKVIMGFPAPKPRNIEKDVKVFPW
XP_755658.      LVGVRFTVEEKNRIRRNLEGFRPLTVSKAKADSEDFFKVIMGFPAPKPRNIEKDVKVFPW
g19421.t1       LVDANFTVEEKNRIRRNLEGFRPITVSKVKSECEEFFKVIMAFPSPKPRNIEKDVKIFHW
XP_0072880      LLNSQFSVEEKNRIRRNLEGFHPMTVSKGKADSEEFFKVIMAFPAPKPRNIEKDVKVFHW
XP_0080841      LVAARFTVEEKNRIRRNLEGYRPLTVSKGKPDSEEFFKVIMGFPTPKPRNIEKDVKVFHW
APA12674.1      LVAAKFTVEEKNRIRRNLEGFRPATVSKAKPDSEEFFKVIMAFPNPKPRNIEKDVKVFHW
XP_0245519      LVAAKFTVEEKNRIRRNLEGFRPATVSKGKPDSEEFFKVIMAFPNPKPRNIEKDVKVFHW
CZT12971.1      LVAGRFTVEEKNRIRRNLEGFHPVTVSKGKPDSEEFFKIIMAFPAPKPRNIEKDVKVFKW
CZT42388.1      LVAGRFTVEEKNRIRRNLEGFHPVTVSKGKSDSEEFFKIIMAFPAPKPRNIEKDVKVFKW
KIN00506.1      LVAARFTVEEKNRIRRNLEGFRPLTVSKGKAESEEFFKVIMGFPTPKPRNIEKDVKVFHW
XP_0310090      LVAVQFTVEEKNRIRRNLEGFHPVTVSKGKSDSEEFFKVIMAFPTPKPRNIEKDVKVFHW
BLGH_06013      LVAAQFTVEEKNRIRRNLEGFRPLTVSKGKSDSEEFFKVIMAFPNPKPRNIEKDVKVFHW
EPQ64905.1      LVAAQFTVEEKNRIRRNLEGFRPLTVSKGKSDSEEFFKVIMAFPNPKPRNIEKDVKVFHW
POS86580.1      LVAAQFTVEEKNRIRRNLEGFRPLTVSKGKPESEQFFKVIMAFPTPKPRNIEKDVKVFHW
RKF58170.1      LVAAQFTVEEKNRIRRNLEGFRPLTVSKGKSESEEFFKVIMAFPVPKPRNIEKDVKVFQW
                *:  .*:*************:.* **** *.:.*:***:**.** ***********:* *

RAQ47439.1      RILGHALKKIIGKYSASYSST-----AGALPTPISST--YTSNGAASDS---GTEPPNAA
XP_755658.      KILSHALKKIIGKYSASYSST-----AGALPTPISST--FAGNGAASDS---GTEPYAAA
g19421.t1       SDLSSALKKIIGKYSASPSSTVAS--ARSPLTQITTNG-FTAERSLIGI-PHSNNHNGPT
XP_0072880      KDLLLTFAQ-----SASPASILPP-----ALTPVSSSTGYGTEGSSTGY--------GGI
XP_0080841      KDLAAALKKIIGKYSASPSSTLPP--APPLLTPSSSTG-YAAEGSSAGI-SYMGDHHGTM
APA12674.1      KDLGSALKKIISKYSASPSSTLPPAHAPALLTPVSSTG-YATETSSVGL-PYTSDHHGGI
XP_0245519      KDLGPALKKIISKYSASPASTLPPTHAPALLTPVSSTG-YATEASSAGL-PYTSDHHGGI
CZT12971.1      KDLTAALKKIISKYSASPSSTLPP--APTLLTPVSSSGDYVTEGSNAAISSYTNDHHGGV
CZT42388.1      KDLTAALKKIISKYSASPSSTLPP--APTLLTPVSSSGDYVTEGSNAGISSYRNDHHGGV
KIN00506.1      KDLASALKKIISKYSASPSSTLPP--APALLTPVSSTG-YGTEGS---------------
XP_0310090      KDLASALKKIFGKYSAGPASTLPP--APALLTPVSSTG-YATEGSSAGL-SYASDHHGAI
BLGH_06013      HDLCGALKKIIGKYSASPSSTVPP--VPPLLTTANLNG-YANNSSTSSL--YSHEHNRAT
EPQ64905.1      HDLCGALKKIIGKYSASPSSTVPP--VPPLLTTANLNG-YANNSSSSNL--YAHDNNRTT
POS86580.1      SDLSSALQKIIGKYSASPSSILNT--ASSLLTPISTSG-YTTESSSAPS--CSNDHTRTS
RKF58170.1      KDLCSALKKIIGKYSASPSSTIAP--STMLLTPISTCG-YATESSSCLS--YSNDHNRTS
                  *  :: :     **. :*            *  .     :  : :

RAQ47439.1      SPQSVSDTGPSTTYGAHVAAAAYSPPAHHHPAHSM---------TGTHELRAVL-PAVSQ
XP_755658.      SPQSVSDAAPSSSYCHSIATPAYSPPTGSSGPHMT----------GTVELRAVL-PAVTQ
g19421.t1       SPHSLSSSVISHS-----PSCDILSPLSNDKSKSTPTLIHHYPTNGIPDTRI---PTHNI
XP_0072880      SPRSMANSTTST-YTSNLPA-RVVSPHEHKSL-ML--------HGGPPDLRVGISPHGPE
XP_0080841      SPRSISGSTTSTAYATSIPA-RVLSPHSQKSM-VL--------GPGPSDLRVSHDPHWQG
APA12674.1      SPRSISGSTTSTAYNSNIPAQRAISPQSQKSI-NF--------QGGPPDLRIAV-PHGTQ
XP_0245519      SPRSISGSTTSTSYNSNIPAQRAISPQSQKSI-GY--------QGGPPDLRIAV-PHGAQ
CZT12971.1      SPRSISGSTTST-YTNNMPA-RGLSTHDQKSM-AL--------QGGPPDHRM---PQHPH
CZT42388.1      SPRSISGSTTST-YTNNMPA-RGLSTHDQKSM-AL--------QGGPPDHRM---PQHPH
KIN00506.1      -----------------LPG-RVLSPHLQKSLAAL--------QSGPPDLRVAI-PH-PH
XP_0310090      SPRSITGSTSSTAYAGNMPT-RVLSPHSQKSM-AL--------QGGPPDLRVSL-PQNPH
BLGH_06013      PP-SLPNTITSTAY----PI-RVLSPNNHEKALSMPLSIMQ--SGSHSDLRMAHMPSVQE
EPQ64905.1      PP-SLPNTVTSTGY----PI-RVLSPNNHEKALSMPLSVIQ--SGSHSDLRMAHMSSVHD
POS86580.1      SPRSYSSSTTSTTA----PSIRLMSPPNNEKANCMTVPMF---RGGGADLQLF--PDIIQ
RKF58170.1      SPQSFASSATSTSA----ASLRLMSPQNQEKVVLMPIPPNQFRGNNSPDMRMQFMTDSIN
                                 .     ..                     .  : .     .

RAQ47439.1      ---PYHPMAAPYSYPAVCQQQGDLGLSA-----PVSR------TWEINP-----LINAPG
XP_755658.      ---PYHSMASGYAYPTVCQQQGQLSLTA-----PVSR-----PSWNPNS-----LAHQPS
g19421.t1       ------YGENTSHVPQ-----Y--------------------------------------
XP_0072880      -ASSGWPGGPPHQLPA--HSSYAPLGHQ-----PPPR-----DAWGMYL------ENSPA
XP_0080841      ----GQQQQQQQHMQT--SHQY-AHLTP-----QTAR-----GSWDMAS------YNHAN
APA12674.1      -ESSHWPQGTTHHMPT-PQQHYQAQLGNAAAV-ASAR-----GSWDLSN----YLENSPA
XP_0245519      -ESSHWPQGTTHHMPT-PQQQYQAQLGNAAAVAASAR-----SSWDLSN----YLENGSA
CZT12971.1      -EGSSYWQGGQHHMQG--SHQY-QSLGN-----HSNR-----GTWDMLG------QTGPT
CZT42388.1      -EGSSYWQGGQHHMQG--SHQY-QSLGN-----HSNR-----GTWDMHG------QTGPA
KIN00506.1      -DAAGPWGAAAHHMSA--QASYHAGAGSQAAAQNHNHHQQRAASWDMAS------YLDAS
XP_0310090      -ETPGHWPGGHHHMQA-SQQQYQQHLGS-----QSGR-----ASWDISSVGVAYLDNGPA
BLGH_06013      TNNPNQWQNTHQHMPS--GQIQVQRSQN-----LYPR-----SPWEMEN------YLDPN
EPQ64905.1      TNNPNQWQNAPQHMPS--GQMQVQRSQN-----IYPR-----SSWDMGN------YLDPN
POS86580.1      ----SPWQNSPHRF-------------------QPTR-----------------------
RKF58170.1      -QNP--WYSSLQR-----------------------------------------------


RAQ47439.1      TNGGPT------------------------------------------------------
XP_755658.      ASAAPS------------------------------------------------------
g19421.t1       ------------------------------------------------------------
XP_0072880      TAASQPGSGFGGRRSDGAASVVESGVESVVVAAAAAAAAAAVAATTPTTTTFTWSADAAY
XP_0080841      QAASLTG-----------------------------------------------------
APA12674.1      TAAGTSGSG---------------------------------------------------
XP_0245519      TAAGTSGTG---------------------------------------------------
CZT12971.1      TAGGTSAPL---------------------------------------------------
CZT42388.1      TAGGTSAPL---------------------------------------------------
KIN00506.1      SAATAGGS----------------------------------------------------
XP_0310090      TTAGSSAGS---------------------------------------------------
BLGH_06013      SYVLPHISG---------------------------------------------------
EPQ64905.1      SYVLPHISG---------------------------------------------------
POS86580.1      ------------------------------------------------------------
RKF58170.1      ------------------------------------------------------------


RAQ47439.1      ------------------------------------------------------------
XP_755658.      ------------------------------------------------------------
g19421.t1       ------------------------------------------------------------
XP_0072880      VRDSCRLEIGDEGRGLAWPGLGWPEPDPGRRSSPHMRYASTHLASTRLTWTRLASTHRPV
XP_0080841      ------------------------------------------------------------
APA12674.1      -----------------------------------------------------VDYQGQGQ
XP_0245519      -----------------------------------------------------VDYQGQG-
CZT12971.1      -----------------------------------------------------QSLN----
CZT42388.1      -----------------------------------------------------QSLN----
KIN00506.1      ------------------------------------------------------------
XP_0310090      ------------------------------------------------------------
BLGH_06013      -----------------------------------------------------IA------
EPQ64905.1      -----------------------------------------------------IA------
POS86580.1      ------------------------------------------------------------
RKF58170.1      ------------------------------------------------------------


RAQ47439.1      YNYLTPMSYSVQDPSH--------------------------
XP_755658.      YSYLAPLPYSLPDPSHGC------------------------
g19421.t1       RFQNGPIPQCSGNDF---------------------------
XP_0072880      HPSIYPSIDPSIDPSIHPSTHPLIHPSHLTLSLLSDLPHIIT
XP_0080841      QLANYPRNNNIADNAVSGADNQLVRSQSHQMSLSRT------
APA12674.1      QQHQTPRNVVDASMAGGGNGSRTLSSLQHQQQSQQIPRT---
XP_0245519      QQHQTPRNVVDGSMTGGGNGNRALSSLQHQQQSQQIPRT---
CZT12971.1      LNYSNHRNNPADSAAATGADNRIVRTISAQQQQRT-------
CZT42388.1      LNYSNHRNNPADSAAATGGDNRIVRTISAQQQQQRT------
KIN00506.1      ------------------------------------------
XP_0310090      QSLNYSRSVNVGEQTSDNRIARSLSVQQQSHQMPRT------
BLGH_06013      NIGNPPMPAYTTSKNVVESNQPDPRPLSLTLKMPRS------
EPQ64905.1      NIGNPPIPAYTTSKNVVESNQSDPRPLSLTLKMPRS------
POS86580.1      ---ALLRTQCDYDNQLLGSG----------------------
RKF58170.1      ------------------------------------------
```
